# Defining the cardiac fibroblast secretome in the healthy and infarcted mouse heart

**DOI:** 10.1101/2024.08.06.606146

**Authors:** Jasmin Bahr, Gereon Poschmann, Andreas Jungmann, Martin Busch, Zhaoping Ding, Ria Zalfen, Julia Steinhausen, Thorsten Wachtmeister, Daniel Rickert, Tobias Lautwein, Christina Alter, Junedh M. Amrute, Kory J. Lavine, Karl Köhrer, Patrick Most, Kai Stühler, Julia Hesse, Jürgen Schrader

**Affiliations:** Department of Molecular Cardiology, Medical Faculty and University Hospital Düsseldorf, Heinrich Heine University Düsseldorf, 40225 Düsseldorf, Germany; Institute for Molecular Medicine, Proteome Research, Medical Faculty and University Hospital Düsseldorf, Heinrich Heine University Düsseldorf, 40225 Düsseldorf, Germany; Division of Molecular and Translational Cardiology, Department of Internal Medicine III, Heidelberg University Hospital, 69120 Heidelberg, Germany; Genomics & Transcriptomics Laboratory, Biological and Medical Research Centre (BMFZ), Heinrich Heine University Düsseldorf, 40225 Düsseldorf, Germany; Center for Cardiovascular Research, Department of Medicine, Cardiovascular Division, Washington University School of Medicine, St. Louis, MO 63110, USA; Molecular Proteomics Laboratory, Biological and Medical Research Centre (BMFZ), Heinrich Heine University Düsseldorf, 40225 Düsseldorf, Germany; CARID, Cardiovascular Research Institute Düsseldorf, Medical Faculty and University Hospital Düsseldorf, Heinrich Heine University Düsseldorf, 40225 Düsseldorf, Germany

## Abstract

Cardiac fibroblasts (CF) are key players after myocardial infarction (MI), but their signaling is only incompletely understood. Here we report a first secretome atlas of CF in control (cCF) and post-MI hearts (miCF), combining a rapid cell isolation technique with SILAC and click chemistry. In CF, numerous paracrine factors involved in immune homeostasis were identified. Comparing secretome, transcriptome (SLAMseq), and cellular proteome disclosed protein turnover. In miCF at day 5 post-MI, significantly upregulated proteins included SLIT2, FN1, and CRLF1 in mouse and human samples. Comparing the miCF secretome at day 3 and 5 post-MI showed the dynamic nature of protein secretion. Specific in-vivo labeling of miCF proteins via biotin ligase TurboID using the POSTN promotor mirrored the in-vitro data. In summary, we have identified numerous paracrine factors specifically secreted from CF in mice and humans. This secretome atlas may lead to new biomarkers and/or therapeutic targets for the activated CF.

## Introduction

Cardiac fibrosis is a common feature of ischemic heart disease and several other disease states, such as diabetes mellitus, and aging^1,2^. Fibrosis increases myocardial stiffness, thereby impairing cardiac function, which ultimately progresses to end-stage heart failure. Numerous studies emphasized, that the severity of cardiac fibrosis in patients correlates with adverse cardiac events and mortality^3^. However, no treatment is presently available to prevent excessive fibrosis.

Cardiac fibrosis is defined as an increase in myocardial extracellular matrix (ECM) deposition by cardiac fibroblasts (CF)^4^, in that the equilibrium between synthesis and degradation of the individual ECM components is disturbed. Proteomic analysis revealed that 90% of cardiac ECM is composed of 10 different proteins, including collagens (collagens I, III, and IV), non-collagenous glycoproteins (fibronectin and laminins), proteoglycans, glycosaminoglycans, and elastins^5^. A fundamental feature of the ECM is that it is altered during inflammation, injury, or infection and with age^6^. It is now being appreciated, that CF not only secrete ECM components, that provide dynamic tissue organization and integrity, but also signaling molecules participating in many biological functions^7^. The endogenous mechanisms that restrain pro-fibrotic signals to protect the myocardium from progressive fibrosis are presently unknown.

Recent single-cell transcriptomics data suggest that CF express various paracrine factors which may signal to surrounding cells^8,9^. To this end, multiple secretory pathways (classical exocytosis and unconventional protein secretion (UPS)) have evolved for optimal precision and timing of intercellular dialogues^10^. Despite its importance in cell-cell interaction, the individual proteins which are secreted from MI-activated CF so far have neither been fully defined nor has their communication with surrounding cells been entirely explored.

In previous studies, the release of bioactive factors from CF has been assessed mostly indirectly and documented the profound effects of CF-conditioned medium on cardiomyocytes^11,12^. Interestingly, mice ablated for PDGFRα^+^ fibroblasts better preserved cardiac function after MI as compared to wildtype controls^13^, again suggesting that the secretome of CF is functionally important. However, up to now there is no study that has rigorously assessed the secretome of CF under in-vivo like conditions. All previous studies reported on the secretome of CF isolated from mice^14^ and from human endomyocardial biopsies^15^ were carried out under serum-free conditions, which very likely critically influence the cellular secretome^16^.

The present study is the first to define the CF secretome in the non-ischemic and the ischemic heart using a combination of a rapid CF isolation technique together with stable isotope labeling using amino acids (SILAC) and click chemistry-based protein enrichment which permitted analysis under optimal, i.e. serum-containing, conditions. Aside of many proteins of the ECM, we identified numerous paracrine/autocrine factors that are specifically secreted from CF of healthy and infarcted hearts in mice and humans. The identified proteins in this secretome atlas may serve as a rich source to study in the future in more detail cell-cell interactions and/or to identify novel diagnostic markers for the infarct-activated fibroblast.

## Results

### Secretome analysis of cCF

For the comprehensive secretome analysis of CF isolated from the unstressed control heart (cCF) and the post-MI heart (miCF), we used a fast (9 min) cell isolation protocol^17^ and short-term culture with 10% FBS, combined with click chemistry-based protein enrichment and SILAC labeling for robust protein quantification by LC-MS/MS^18^ (Figure 1).

**Figure 1:**
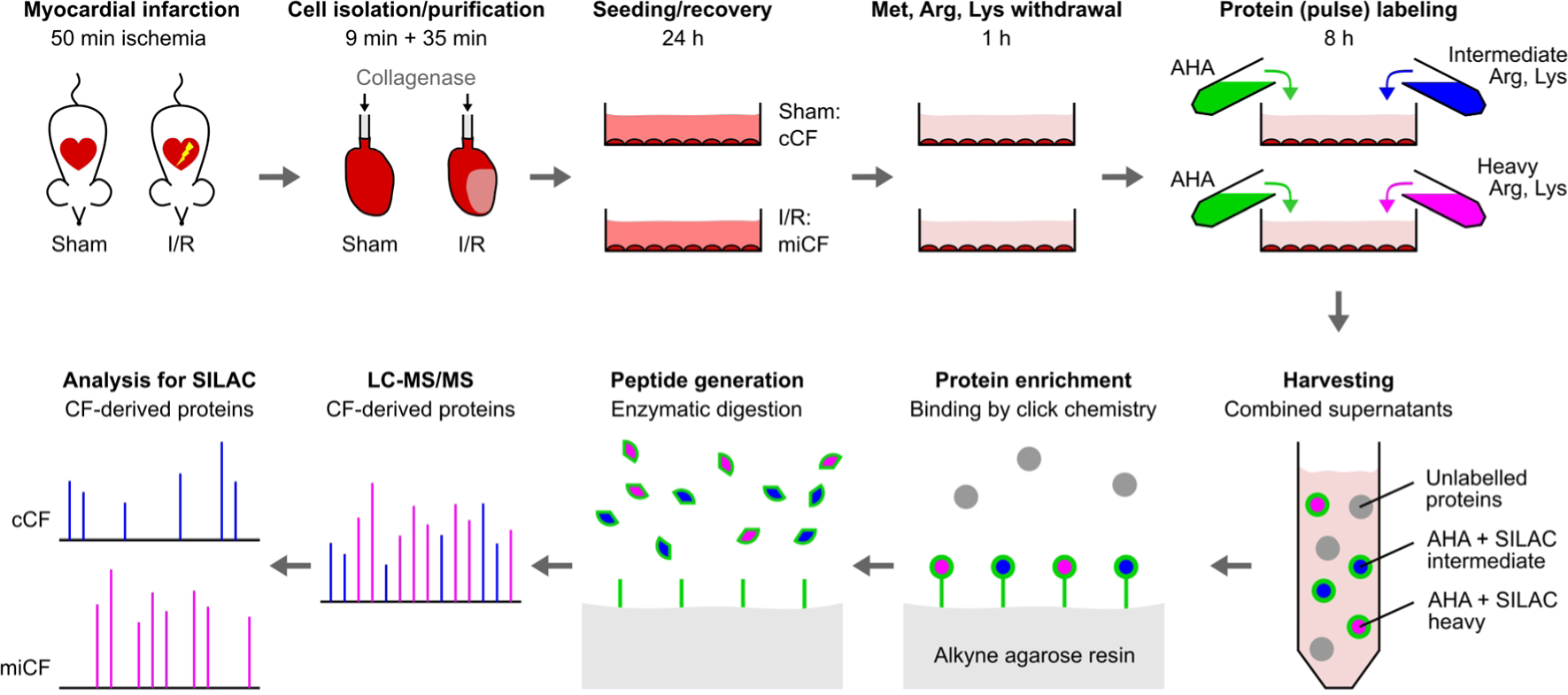
Workflow of comparative secretome analysis of CF from sham control hearts (cCF) and infarcted hearts (miCF). CF were isolated from mouse hearts 5 days after MI (50 min ischemia/reperfusion, I/R; miCF) or sham surgery (sham; cCF) and cultured in conventional cell culture medium containing 10% FBS for 24 h. Amino acids L-methionine (Met), L-arginine (Arg), and L-lysine (Lys) were withdrawn for 1 h and subsequently replaced by azidohomoalanine (AHA) together with either intermediate or heavy Arg and Lys isotypes ([^13^C6] Arg, [4,4,5,5-D4] Lys or [^13^C6, ^15^N4] Arg, [^13^C6, ^15^N2] Lys, respectively). After incubation for 8 h, cCF and miCF supernatants were harvested and combined. Newly synthesized, AHA-containing proteins were bound to an agarose-resin by click chemistry. After stringent washing to remove non-bound proteins, remaining proteins were enzymatically digested and eluted peptides were applied to LC-MS/MS. In the following data analysis, intermediate and heavy isotype labels were used to assign peptides to cCF and miCF samples.

In cCF isolated from sham control hearts we identified a total of 122 proteins, of which the majority of proteins (92%) were predicted to be secreted (Figure 2A), either conventionally (Conv) or via unconventional protein secretion (UPS), indicating active release from cCF. In line with the key role of CF in maintaining structural tissue integrity, ECM-associated proteins accounted for about two thirds of the total protein amount (Figure 2B). A major fraction of ECM proteins were collagen chains (Figure 2C). We also found substantial secretion of proteins involved in collagen stabilization or turnover, such as lysyl oxidases (LOX, LOXL1-3), lysyl hydroxylases (PLOD1, 3), matrix metalloproteinases (MMP2, 3), as well as tissue inhibitors of metalloproteinases (TIMP1-3), which together are likely to be involved in continuous matrix maintenance.

**Figure 2:**
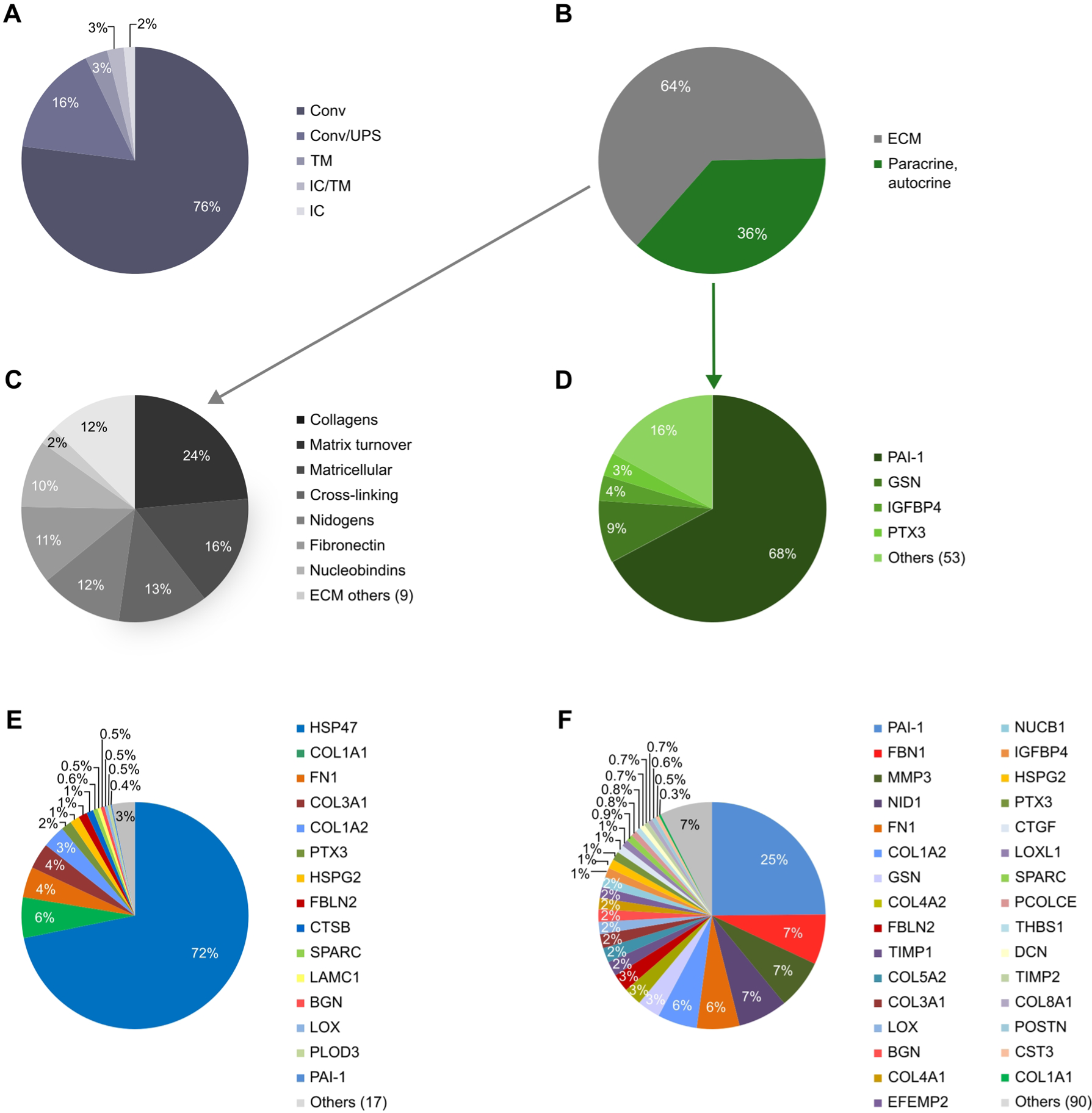
Secretome analysis in cCF. LC-MS/MS data identified 122 secreted proteins in cCF isolated from sham control mouse hearts 5 days after surgery (n=4; source data in Supplemental Data 1). **A)** Protein localization as predicted by OutCyte57. Shown is the percentage of released proteins classified as conventionally secreted (Conv), unconventionally secreted (UPS), intracellular (IC), and transmembrane (TM) proteins. **B)** Protein intensity fractions of cCF secretome categorized as ECM-associated proteins and paracrine/autocrine factors according to KEGG annotation and PubMed search. **C)** Subcategories of ECM-associated proteins. **D)** Subcategories of proteins assigned as paracrine/autocrine factors. **E)** Relative intracellular levels (intensities) measured in cell lysates of proteins that were identified in cCF secretome analysis (32 of the 122 secreted proteins). **F)** Protein intensity representation of all proteins secreted from cCF.

In addition to the ECM-associated proteins, the cCF secretome contained multiple proteins with potential paracrine and/or autocrine function (Figure 2D), which amounted to 36% of the total measured protein intensity (Figure 2B). The most abundant paracrine factor was plasminogen activator 1 (PAI-1, *Serpine1*), representing 68% of paracrine/autocrine protein intensity (Figure 2D) and 25 % of the total secretome (Figure 2F). PAI-1 has been shown to control cardiac TGF-β production and loss of PAI-1 results in cardiac fibrosis^19^. Supplemental Table 1 summarizes secreted proteins with reported biological function. In addition to PAI-1, this includes gelsolin (GSN), secreted frizzled-related protein-1 (SFRP1) and slit homolog 2 protein (SLIT2) as well as proteins influencing the complement system such as pentraxin-related protein 3 (PTX3) and complement C3 (C3). Immune homeostasis may be influenced by tyrosine-protein kinase receptor UFO (AXL), chemokine (C-C motif) ligand 2 (CCL2) and superoxide dismutase 3 (SOD3). Angiogenesis may be promoted by insulin-like growth factor binding protein 4 (IGFBP4), angiopoietin-like 4 (ANGPTL4), beta-nerve growth factor (NGF), and pigment epithelium-derived factor (PEDF, *Serpinf1*). Finally, proteins were secreted that are known to confer cardioprotection such as IGFBP4, NGF, and SFRP1, as well as proprotein convertase subtilis/kexin 6 (PCSK6). Note that several of the above proteins have previously not been assigned to CF as a primary site of production, such as AXL, IGFBP4, and PCSK6. Together these data suggest that cCF secrete numerous biologically active proteins that can signal to a variety of surrounding cardiac cells to maintain tissue integrity and homeostasis under unstressed conditions.

To gain further insight into the dynamics of protein secretion/turnover from cCF, we compared our secretome data with corresponding intracellular proteome data (Figure 2E) and transcriptome data that were generated by SLAMseq (see Methods). The graphical display of combined data in Figure 3 enables a direct comparison of the secretome, proteome and transcriptome for all proteins secreted by cCF. It can be seen that PAI-1 (*Serpine1*), in contrast to its high secretion (Figure 2F), amounted to only 0.4% of the intracellular intensity of secreted proteins (Figure 2E). Together with substantial PAI-1 transcript levels and a high T>C conversion rate indicating active RNA synthesis (Supplemental Table 2), this suggests rapid intracellular turnover and immediate secretion of this central suppressor of cardiac fibrosis. In contrast, heat shock protein 47 (HSP47, *Serpinh1*), which is a member of the serpin protein family involved in assembly of triple-helical procollagen molecules in the ER^20^, accounted for 72% of the intracellular intensity of secreted proteins (Figure 2E) and showed substantial transcription, but only low secretion (Figure 3, Supplemental Table 2). Similarly, collagen chains (COL1A1, COL1A2, COL3A1) showed high intracellular protein pool sizes and high mRNA transcript levels, but comparatively little secretion (Figure 3). This might suggest that collagens are only secreted when properly assembled. Also note, that many of the proteins secreted from cCF were below detection limit in the intracellular proteome analysis (90 of 122 proteins), indicating small intracellular protein pool sizes and therefore high cellular turnover. This includes paracrine factors, such as insulin-like growth factor (IGF1) and the chemokine CCL2, also known as monocyte chemoattractant protein 1 (MCP1) (Supplemental Table 2).

**Figure 3:**
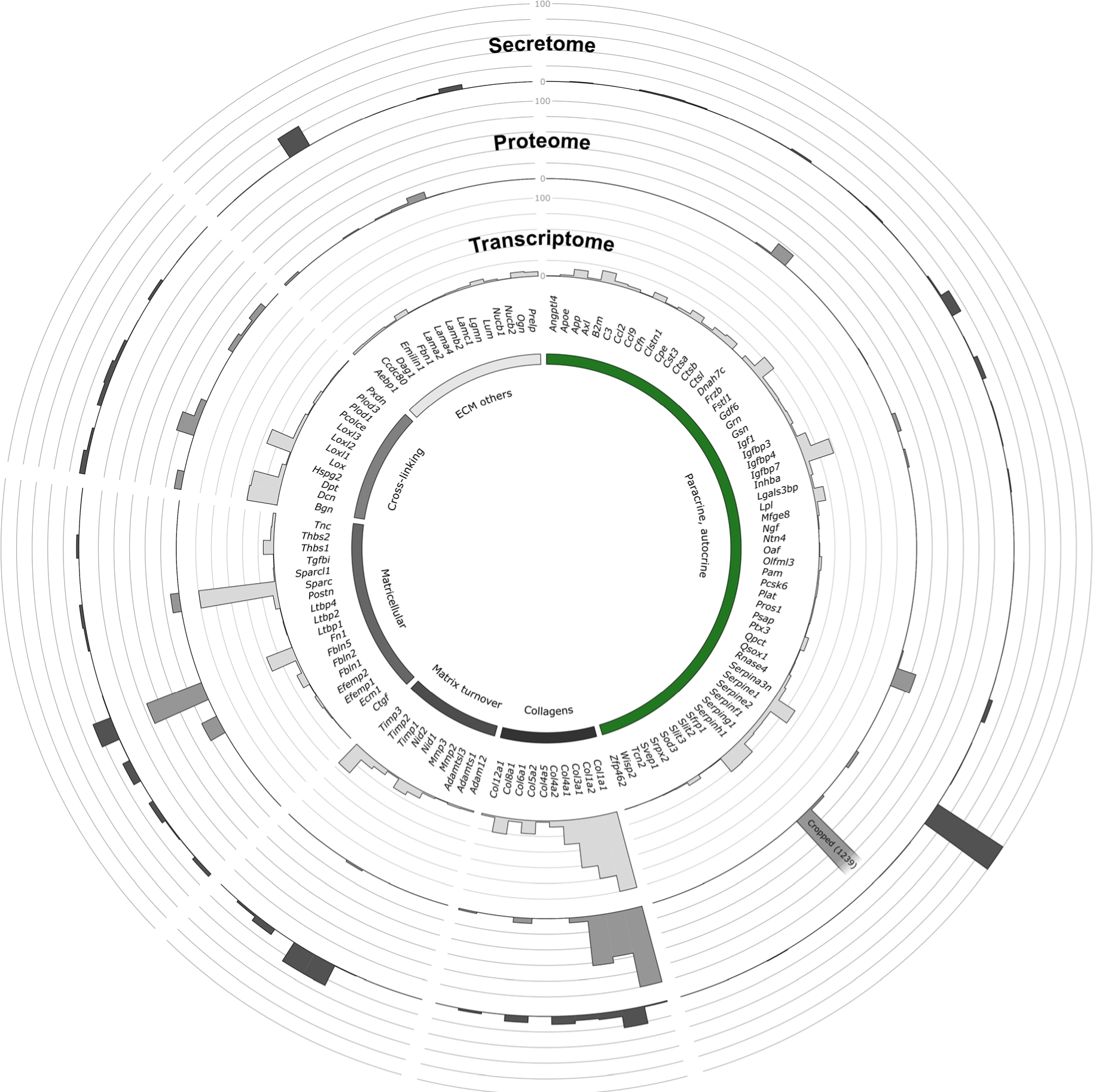
Comparison of secretome, proteome, and transcriptome in cCF. Gene expression (transcript levels, light grey bars; source data in Supplemental Data 3), cellular proteome (protein intensities, grey bars; source data in Supplemental Data 2), and secretome (protein intensities, dark grey bars; source data in Supplemental Data 1) of the 122 proteins identified in cCF secretome analysis. Data are shown in a Circos68 plot as means of n=3 (transcriptome) and n=4 (secretome, proteome) of cCF preparations from unstressed hearts. To permit comparison, mean value of the gene/protein with the highest expression in the transcriptome (COL1A1, *Col1a1*) and secretome (PAI-1, *Serpine1*) was set to 100%. Due to the dominance of HSP47 (*Serpinh1*) in the proteome, the second highest value (COL1A1) was set to 100% and HSP47 (=1239%) was cropped to allow visualization of lower abundant proteins.

To address cell-cell-communication between secreted paracrine factors and target receptors on individual cardiac cell types, receptors for paracrine factors were selected from the ligand-receptor database of CellTalkDB^21^ and their expression was analyzed in a publicly available snRNAseq data set of murine cardiac cells including CM^22^. The interactome plot (Supplemental Figure 2) revealed, that cCF-derived paracrine factors can signal to multiple cardiac cell types because of the broad expression of their respective target receptors.

To explore whether the secreted proteins are CF-specific, we searched a published murine cardiac scRNAseq data set^22^ for the cellular distribution of the identified secretome proteins. As summarized in Supplemental Figures 3 and 4, many of the proteins were found to be predominantly expressed by CF. Particularly GSN, highly secreted from cCF (Figure 2D), showed high expression levels (Supplemental Figure 4). Also note, that C3 and SLIT3 were highly and preferentially expressed in epicardial cells. To further explore the translational potential of our findings in mice, we made use of recent snRNAseq and CITE-seq data sets generated by two of the coauthors (K.L. and A.J.) on human donor hearts, which were obtained from brain-dead individuals with no known cardiac disease and normal LV function^23–25^. A direct comparison of the mouse with human data showed a remarkable similar distribution pattern at the gene expression level of the extracellular matrix proteins (Supplemental Figure 3) and secreted paracrine/autocrine factors (Supplemental Figure 4). Similar to mice, GSN was highly expressed in human CF and both C3 and SLIT3 transcripts were preferentially found in human epicardial cells (Supplemental Figure 4).

### MI-induced changes in the CF secretome

To explore MI-induced changes in the CF secretome, we isolated activated fibroblasts (miCF) from infarcted mouse hearts (50 min I/R, 5 days after surgery). As shown in Figure 4A, we identified 31 proteins which were only detected in the miCF secretome together with 122 proteins that were already present in the cCF secretome. Of all 153 identified proteins, 28 proteins were found to be significantly upregulated when comparing the miCF with the cCF secretome (Figure 4B).

**Figure 4:**
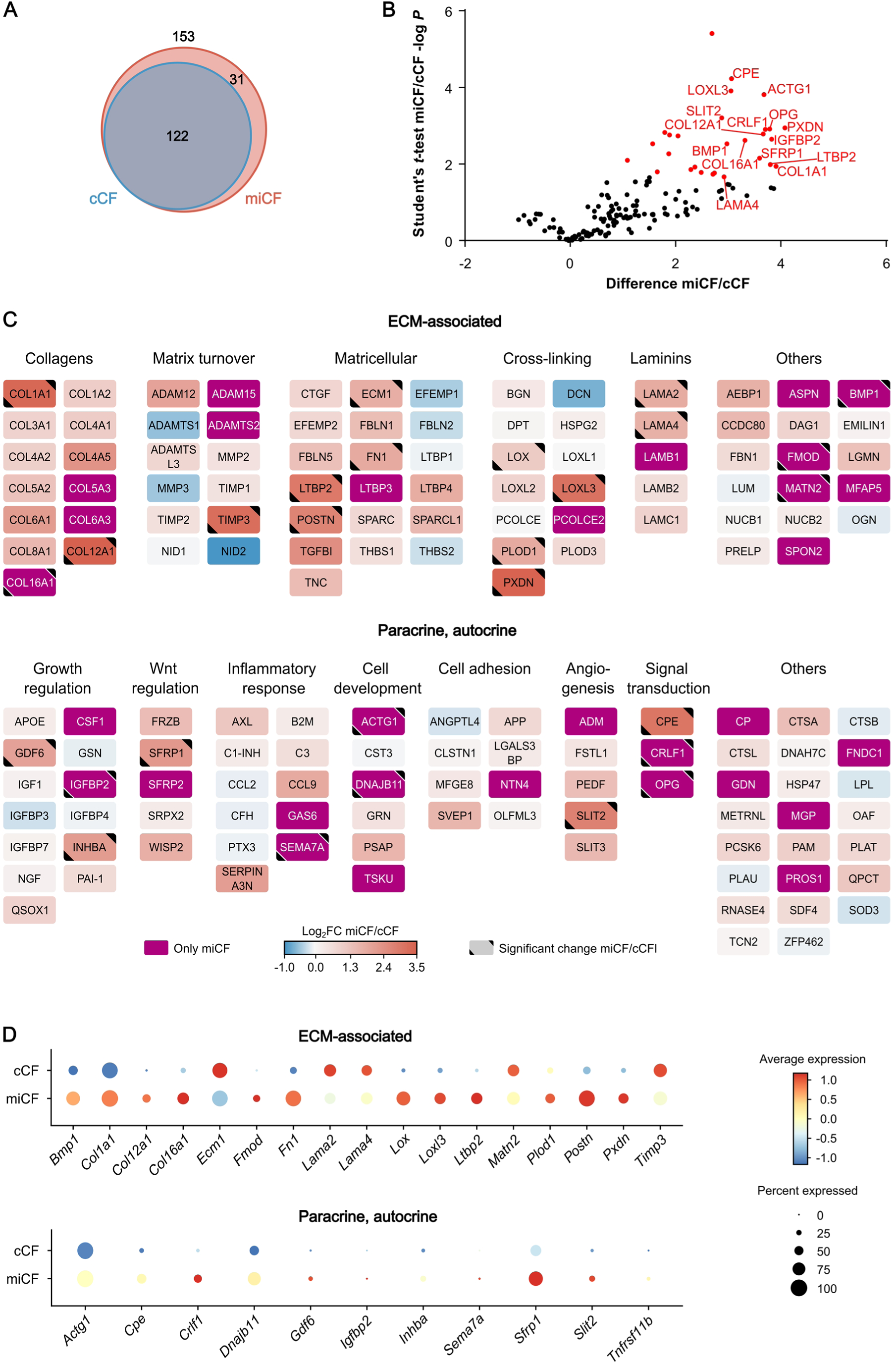
MI-induced changes of the CF secretome 5 days after infarction. LC-MS/MS analysis identified 153 secreted proteins from post-MI CF (miCF) isolated from mouse hearts 5 days after I/R surgery (n=4; source data in Supplemental Data 1). These data were compared to the basal secretome data (122 proteins) of cCF isolated from sham-operated hearts 5 days after surgery (n=4; source data in Supplemental Data 1). A) Venn diagram69 of identified proteins. B) Volcano plot of differentially secreted proteins. Proteins with significantly different intensities in cCF and miCF samples (Student’s *t*-test-based SAM analysis, 5% FDR, S0=0.1) are highlighted in red (28 proteins). For statistical significance analysis of secreted proteins which were only detected in miCF, an imputation approach of missing base values was performed, using values taken from a downshifted normal distribution (details in Methods). Names of top 15 proteins with highest difference between cCF and miCF are annotated. The shown difference refers to the difference of group mean values of log2 transformed intensities. C) Log2 fold changes (FC) of protein intensities between miCF and cCF samples visualized with Cytoscape70. Proteins were grouped in subcategories ECM-associated proteins and paracrine/autocrine factors. D) Transcript levels of the 28 significantly changed proteins in the transcriptome of CF of mouse hearts 5 days after I/R (miCF) or sham surgery (cCF) (n=3 each). Previously published scRNAseq data of cCF and miCF8 were re-analyzed and expression levels in the total cCF/miCF population are visualized as dot plots.

Figure 4C summarizes in a graphical manner the changes in the 153 proteins categorized into ECM-associated proteins and paracrine/autocrine factors. Some cardiac functions reported in the literature for the identified proteins are listed in Table 1. As can be seen, the ECM proteins that were found to be significantly enriched (Figure 4C) included several proteins with known pro-fibrotic function, such as bone morphogenetic protein 1 (BMP1, only detected after MI), lysyl oxidase homolog 3 (LOXL3), periostin (POSTN), extracellular matrix protein 1 (ECM1), but also fibromodulin (FMOD, only detected after MI) with reported strong anti-fibrotic action. Levels of collagen chains COL1A1, COL12A1 and COL16A1 (only detected after MI) were also significantly enhanced (Figure 4C). Increased proteins also included TIMP3, known to promote angiogenesis, as well as peroxidasin (PXDN) and fibronectin (FN1), which both have a reported pro-apoptotic activity in cardiomyocytes^26,27^. Note, that levels of several proteins like ADAM metallopeptidase with thrombospondin type 1 motif 1 (ADAMTS1), MMP3, nidogen-2 (NID2), EGF-containing fibulin-like extracellular matrix protein 1 (EFEMP1), thrombospondin 2 (THBS2), and decorin (DCN) were reduced in the secretome of miCF as compared to cCF (Figure 4C), but differences did not reach the level of significance.

**Table 1:**
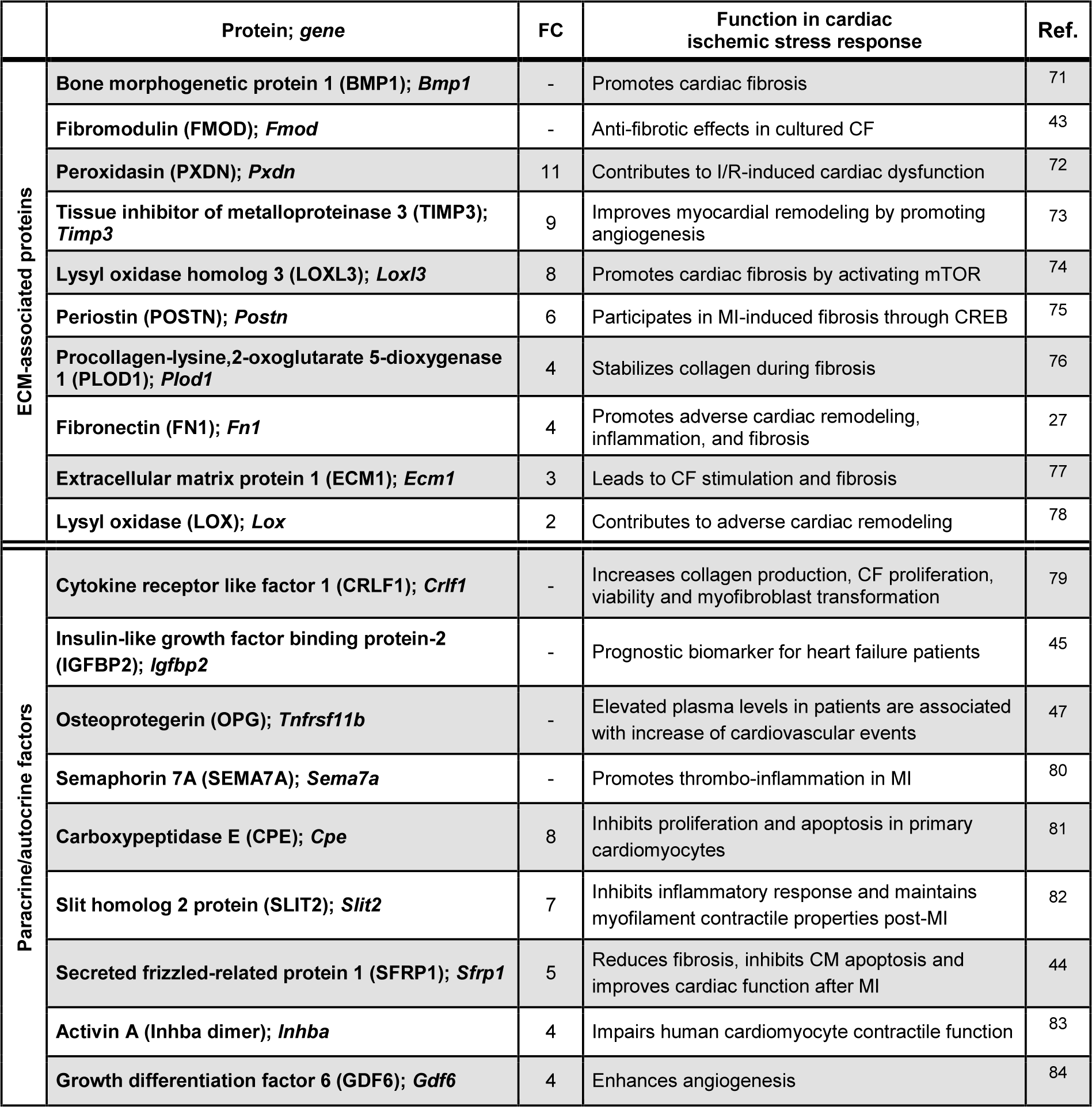
Reported functions of selected ECM-associated proteins and paracrine/autocrine factors significantly enriched in the miCF secretome as compared to the cCF secretome. Proteins with significant changes between the secretomes of CF isolated from the uninjured heart (cCF) and infarcted heart (miCF) 5 days after surgery were assigned and selected by their KEGG annotation and PubMed search. Proteins are listed in decreasing miCF/cCF fold changes (FC) of secretome intensities. Proteins only detectable in miCF are listed at the top. Quantitative information of the listed proteins can be found in Supplemental Data 1.

In addition to the 17 significantly enriched ECM-related proteins, we identified 11 significantly enhanced paracrine/autocrine factors in the miCF secretome (Figure 4C). Among these were semaphorin 7A (SEMA7A; only detected after MI), which is known to mediate thrombo-inflammation, and growth differentiation factor 6 (GDF6), which can promote angiogenesis (Table 1). There were several proteins with reported cardioprotective activity, such as carboxypeptidase E (CPE), Wnt/β-catenin pathway inhibitor SFRP1, and SLIT2. Notably, two proteins only detected in the miCF secretome, insulin-like growth factor binding protein 2 (IGFBP2) and osteoprotegerin (OPG), have already been reported as prognostic biomarker in human heart failure and additional cardiovascular events, respectively. Together these data demonstrate that proteins secreted by miCF are not only involved in post-MI fibrosis and scar formation, but also potentially can signal to surrounding cells such as cardiomyocytes, immune cells and coronary vessels to modulate remodeling processes in the injured tissue.

To study whether the enhanced protein secretion by miCF is reflected by respective changes in gene expression, we analyzed scRNAseq data sets previously obtained by us under identical experimental conditions (CF isolated from sham control hearts and infarcted hearts 5 days post I/R)^8^. As displayed in Figure 4D, 12 out of the 17 enhanced ECM-related secretome proteins and all of the enhanced paracrine factors also showed higher gene expression levels in miCF compared to cCF. This indicates that upregulation at the transcriptional level can explain the increased protein secretion.

To investigate, whether the identified 28 proteins that were significantly enriched in the miCF secretome are also predominantly expressed in CF in comparison to other cardiac cell types, we made use of scRNAseq data previously published by us^28^ as well as data from acute MI patients (AMI, <3 months post-MI)^23–25^. As summarized in Figure 5A, genes encoding the enriched ECM-related secretome proteins and paracrine/autocrine factors were for the most part selectively expressed in miCF in mice and humans. Note, that the expression of the proteins was rather heterogeneously distributed within the different miCF populations in mice. The overlap between mice and humans was less pronounced for paracrine/autocrine factors (Figure 5B).

**Figure 5:**
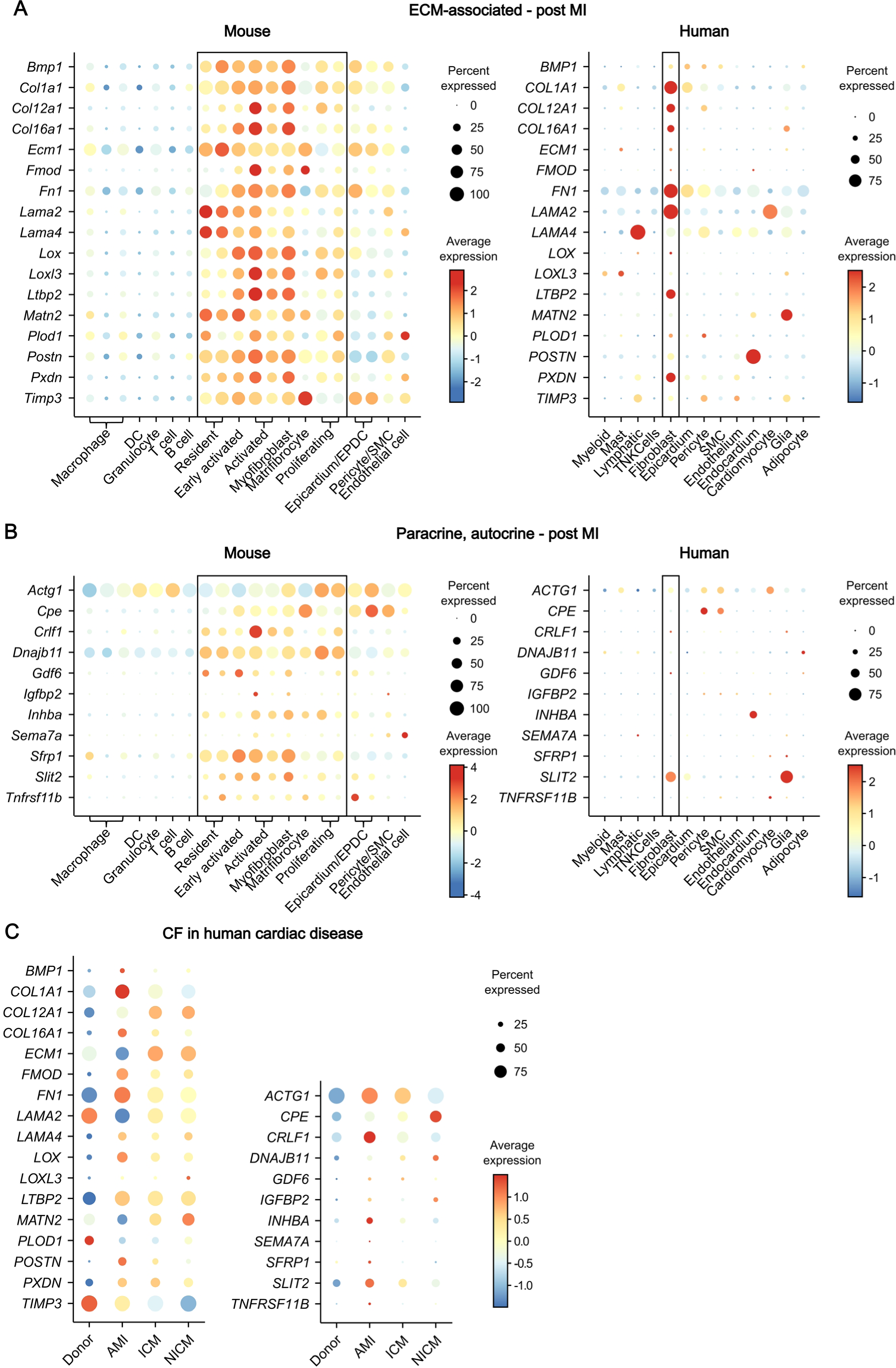
CF specificity of significantly altered secretome proteins post-MI. **A-B)** Transcript levels of the 28 significantly changed proteins between the miCF secretome and the cCF secretome in the transcriptome of cell populations of n=3 infarcted mouse hearts 5 days after I/R (left panels) and of n=3 human heart specimen of acute MI (AMI, <3 months post-MI) patients (right panels). Previously published scRNAseq data of murine stromal and immune cell populations28 and snRNAseq data of human cardiac cells23–25 were re-analyzed. Expression levels are visualized as dot plots, subdivided in **A)** ECM-associated proteins and **B)** paracrine and autocrine factors. Values of CF populations are enframed. DC, dendritic cell; EPDC, epicardium-derived cell; SMC, smooth muscle cell. **C)** To compare the expression of the miCF secretome proteins in CF in human MI and heart failure, we used CITE-seq data25 of human left ventricle (LV) CF obtained from n=6 healthy donors, n=4 acute MI (AMI) patients (<3 months post-MI), n=6 ischemic cardiomyopathy (ICM, >3 months post-MI) patients, and n=6 non-ischemic cardiomyopathy (NICM, idiopathic dilated cardiomyopathy) patients.

The available human data set^23–25^ contains besides CF samples from healthy individuals (donor) and AMI patients also samples from, ischemic cardiomyopathy (ICM, >3 months post-MI) patients and non-ischemic cardiomyopathy (NICM, idiopathic dilated cardiomyopathy) patients. This permitted us to explore whether the identified secreted proteins are infarct-specific or can also be observed in other heart pathologies such as ICM and NICM. As summarized in Figure 5C, gene expression of most of the identified proteins was preferentially upregulated in AMI CF. ICM and NICM CF showed a different signature, with high expression levels of COL1A1, ECM1, matrilin-2 (MATN2), and CPE. Together these data indicate, that the secretome of CF in the human heart is critically determined by the underlying cardiac disease.

To obtain insight into the temporal changes of protein secretion from MI-activated CF we carried out identical experiments as shown in Figure 4 (day 5 post-MI), at day 3 post-MI (Supplemental Figure 5). Comparing both time points, we found overall more proteins to be significantly changed in comparison to sham control on day 3 (80) than at day 5 post-MI (29), with an overlap of 21 proteins (Figure 6A and Supplemental Table 3). Again, numerous ECM-associated proteins and paracrine/autocrine factors were significantly upregulated (Supplemental Figure 5C) of which CCL9, granulin (GRN), LOXL2, osteoglycin (OGN), and SLIT2 were reported in the literature to be involved in fibrosis development. From the temporal changes (Figure 6B) it can be seen that the secretion of some proteins increased over time (POSTN, INHBA, SLIT2) while the others peaked at day 3 and thereafter decreased to a different extent. Particularly, secretion of CPE, COL1A1, and FN1 dramatically decreased after day 3. Together these findings are in line with the notion that CF are a very dynamic cell type that achieves selective differentiated states in the process from acute wound healing to long-term remodeling^29^.

**Figure 6:**
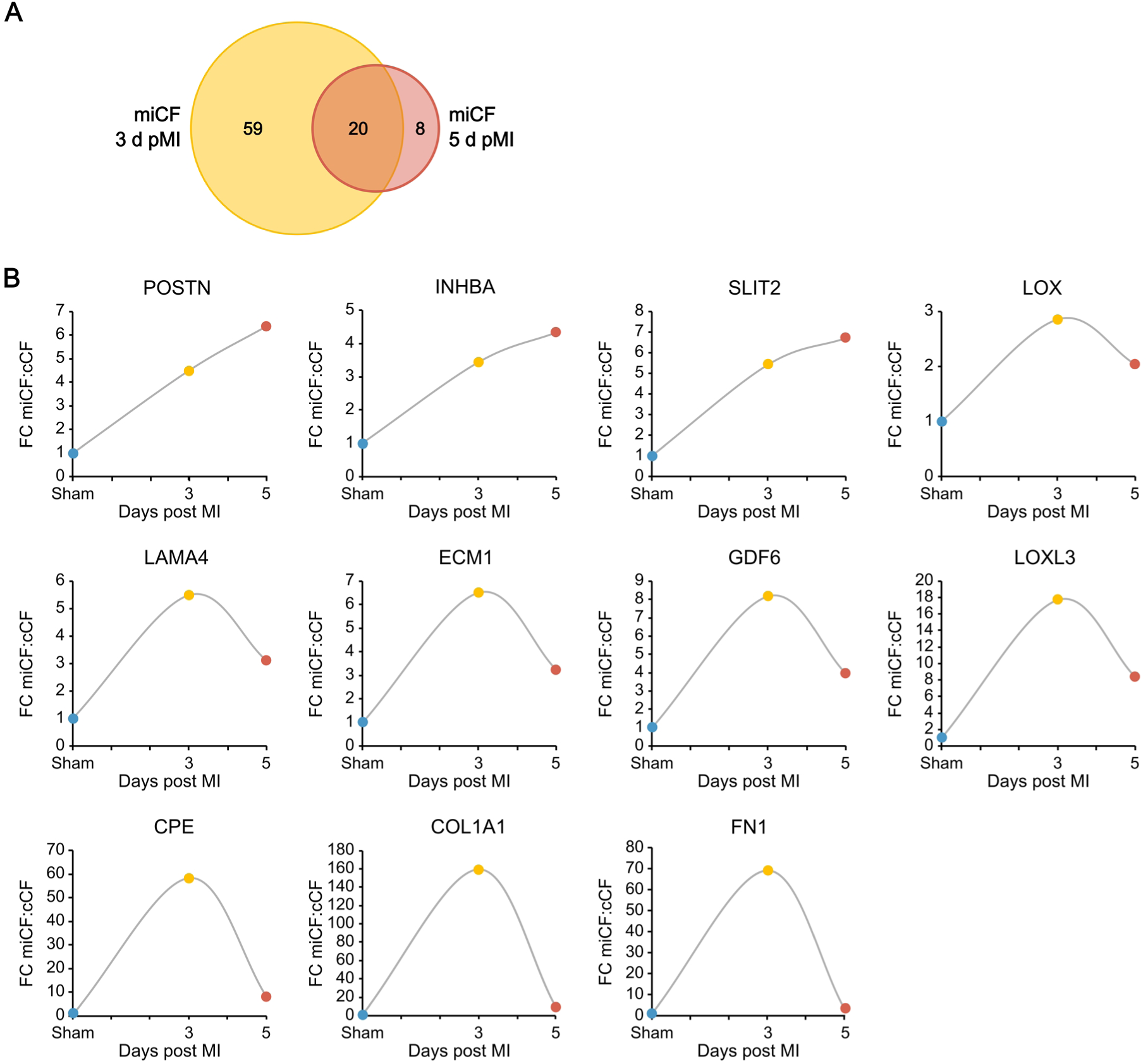
MI-induced changes of the CF secretome 3 and 5 days after infarction. LC-MS/MS analysis identified 129 secreted proteins from post-MI CF (miCF) isolated from mouse hearts 3 days after I/R surgery (n=4; source data in Supplemental Data 1): These data were compared to the secretome data (153 proteins) of miCF from mouse hearts 5 days after I/R surgery (n=4; source data in Supplemental Data 1). **A)** Venn diagram69 of significantly changed secretome proteins of miCF in comparison to the secretome proteins of cCF from sham-operated hearts 3 and 5 days after surgery (n=4 each, source data in Supplemental Data 1), respectively. **B)** Fold changes of selected secretome proteins between miCF and cCF at days 3 and 5 post I/R and sham surgery, respectively.

In a final set of experiments, we made an attempt to validate our in-vitro data under in-vivo conditions. To specifically label proteins that that are released via the ER secretory pathway by activated POSTN^+^ CF, we used the ER-localized biotin ligase ER-TurboID expressed under control of the POSTN promotor that was delivered to miCF via an AAV9 vector system (AAV9-POSTN-ER-TurboID). After i.v. injection of AAV9-POSTN-ER-TurboID at day 1 after MI (50 min ischemia/ reperfusion) and biotin administration via the drinking water, hearts were perfused at day 5 post-MI in the Langendorff mode to capture the secreted biotinylated proteins in the coronary effluent, largely devoid of plasma proteins (workflow in Supplemental Figure 6A). Heart perfusion was necessary because POSTN is also expressed in the liver^30^, resulting in ER-TurboID expression and thus biotinylation of liver-derived plasma proteins. As shown in Figure 7A, we have identified 157 secreted proteins in the coronary effluent, of which 27 overlapped with the secretome proteins of cultured miCF (Supplemental Table 4). Comparing the effluent proteins of mice treated with AAV9-POSTN-ER-TurboID and of non-transduced control mice 5 days post-MI revealed 73 proteins with significantly different protein intensities (Figure 7B), very likely representing effluent proteins originating from the POSTN^+^ CF. Of these, 10 overlapped with the miCF secretome (Figure 7A, Supplemental Table 4). Of the in total 27 overlapping proteins, 25 proteins were preferentially expressed in CF in the infarcted heart (Figure 7C).

**Figure 7:**
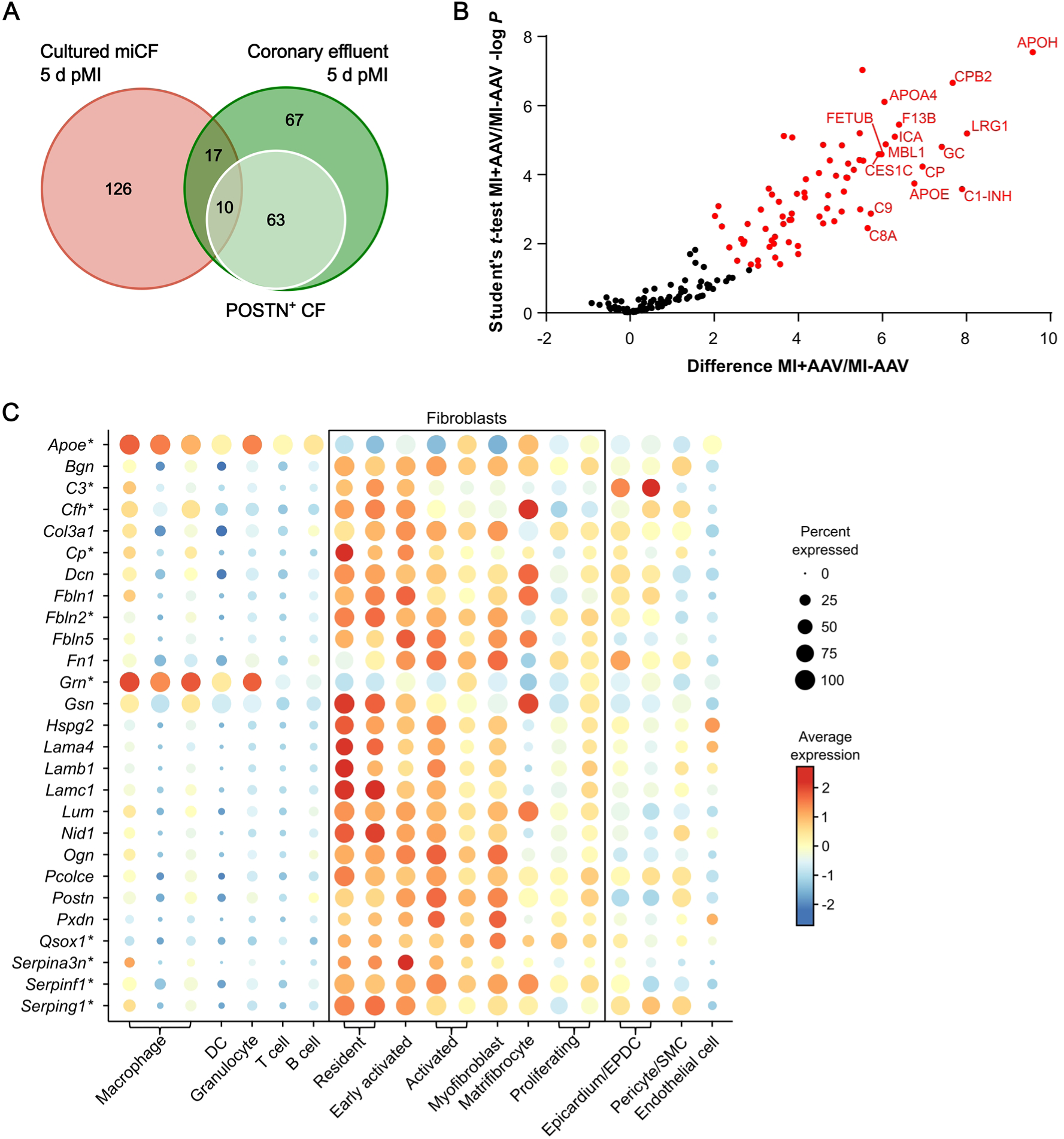
In-vivo-secretome analysis of POSTN+ CF 5 days post-MI. Mice were transduced with AAV9-POSTN-ER-TurboID to allow biotin-labeling and subsequent enrichment of proteins secreted from POSTN+ CFs in the coronary effluent (work flow in Supplemental Figure 6). LC-MS/MS analysis identified 157 proteins in the coronary effluent 5 days after I/R surgery (n=6; source data in Supplemental Data 4). These data were compared to the secretome data (153 proteins) of miCF isolated from hearts 5 days after MI (n=4; source data in Supplemental Data 1). **A)** Venn diagram69 of identified proteins. Green circle, 157 proteins identified in the coronary effluent 5 days post-MI. White circle, 27 proteins significantly enriched in the effluent of AAV9-POSTN-ER-TurboID-transduced mice (n=6) in comparison to non-transduced control mice 5 days post-MI (n=6), representing the in-vivo-secretome of POSTN+ CF. Red circle, 153 proteins in the miCF secretome 5 days post-MI. **B)** Volcano plot of proteins identified in the coronary effluent. Proteins with significantly different intensities in samples from TurboID-transduced mice (MI+AAV) and non-transduced control mice (MI-AAV) (Student’s *t*-test-based SAM analysis, 5% FDR, S0=0.6) are highlighted in red (73 proteins). Names of top 15 proteins with highest difference between MI+AAV and MI+AAV are annotated. The shown difference refers to the difference of group mean values of log2 transformed intensities. **C)** Dot plot visualizing transcript levels of the proteins identified in both data sets in the transcriptome of cell populations of n=3 infarcted mouse hearts 5 days after I/R (left panels). Previously published scRNAseq data of murine stromal and immune cell populations28 were re-analyzed. Genes of proteins that were significantly enriched in the effluent of AAV9-POSTN-ER-TurboID-transduced mice are marked with an asterisk.

## Discussion

This study provides the first comprehensive overview of the secretome of CF in the infarcted heart as compared to the unstressed heart. It demonstrates the fibroblast-specific secretion of numerous paracrine/autocrine factors that can signal to surrounding cardiac cells to potentially influence cardiac remodeling. The newly identified proteins, verified by scRNAseq/snRNAseq in mouse and human tissue samples, may serve as a rich source for novel therapeutic targets as well as biomarkers for CF activation in the context of injury-induced cardiac fibrosis.

A combination of several experimental advances made this study possible. Firstly, we minimized cell activation during the cell isolation procedure by applying a recently developed enzymatic technique which takes only 9 min for tissue digestion^17^ and combined this with short-term culture (2 days) under serum-containing conditions. Secondly, we used click chemistry with AHA together with SILAC which permitted secretome analysis under optimal culture conditions. As a result, the majority (93%) of the identified proteins were predicted to be secreted (Figure 2A). Finally, cell culture data were validated by specific in-vivo labeling of infarct-activated fibroblast proteins via biotin ligase ER-TurboID expressed under control of the POSTN promotor. Together, it appears likely that results obtained under the chosen in-vitro conditions can be extrapolated to the in-vivo situation. This conclusion is further supported by reported scRNAseq/snRNAseq data of mice and humans, showing that most of the genes encoding identified proteins were rather selectively expressed in CF in comparison to other cardiac cell types.

Under homeostatic conditions, the ECM composition is tightly controlled by the equilibrium between synthesis and degradation of its individual components. In line with this concept, we observed that in the secretome of unstressed cCF, collagens accounted for the largest fraction (24%) of ECM-associated proteins, followed by proteins assigned to matrix turnover as the second largest fraction (16%) (Figure 2C). A more direct estimate of the turnover of individual secreted proteins was derived by relating the measured rate of secretion (secretome) to the cellular protein content (proteome) and transcription level (transcriptome) of each protein (Figure 3). Of the proteins secreted under basal conditions (122), the respective intracellular protein was only measurable for a small number of proteins (32). Interestingly, the highest cellular pool sizes were found for FN1, HSP47 (*Serpinh1*), and collagen chains (COL1A1, COL1A2, COL3A1). Note, that among the collagen chains the highest intracellular levels were found for COL1A1, a major protein of the extracellular matrix (ECM), with only little secretion. This is consistent with the reported low turnover of the ECM which was estimated to be of the order of years^31^. It is also remarkable, that the cellular turnover of the secreted proteins varied widely. Very likely it is the imbalance between accumulation of secreted proteins and their rate of degradation/tissue washout, that finally determines the biologically active concentration and leads to tissue fibrosis and cardiac dysfunction.

Of the paracrine/autocrine factors secreted by cCF, PAI-1 comprises with 68% the largest fraction (Figure 2D). PAI-1, a serine protease inhibitor primarily known for regulation of fibrinolysis, is now additionally appreciated to function in many physiological processes including inflammation, wound healing and cell adhesion^32^. Interestingly, global PAI-1 knockout mice develop age-dependent cardiac-selective fibrosis and a similar phenotype was observed in a cardiomyocyte-specific PAI-1 knockout^33^. Thus, both cardiomyocyte- and CF-derived PAI-1 are likely to contribute to cardiac PAI-1 formation and thus may play a role in the preservation of cardiac performance under stress.

Analysis of the cellular distribution of gene expression of secreted proteins in reported scRNAseq/snRNAseq data revealed, that a large fraction of secreted ECM proteins was preferentially expressed in cCF (Supplemental Figure 3). This included various collagen chains, MMP2, fibulin 2 (FBLN2, reported to be essential for angiotensin II-induced myocardial fibrosis^34^), DCN (a proteoglycan possessing powerful antifibrotic, anti-inflammatory, antioxidant, and antiangiogenic properties^35^), fibrillin 1 (FBN1, a constituent of the myocardial ECM in fibrosis^36^), and lumican (LUM, playing an important role in the cardiac fibrillogenesis^37^).

Among the paracrine/autocrine factors, the expression of GSN, PEDF, follistatin-related protein 1 (FSTL1), and AXL was predominantly found in cCF (Supplemental Figure 4). GSN has been reported to be an important mediator of angiotensin II-induced activation of CF and fibrosis^38^, FSTL1 was shown to induce cardiac angiogenesis^39^, AXL is likely to play a role in target organ inflammation^40^, and PEDF was shown to display a protective role on endothelial tight junctions and the vascular barrier in the MI heart^41^. Together these data demonstrate, that non-activated CF secrete numerous biological active proteins which according to the literature are essential for balancing important biological processes such as angiogenesis, inflammation, and fibrosis in the unstressed heart.

In the secretome of infarct-activated miCF, we identified a total of 28 proteins to be significantly enriched. Besides 17 ECM-associated proteins, this included 11 paracrine factors (Figure 4C), of which nine have a reported cardiovascular functionality (Table 1), such as promotion of thrombo-inflammation (semaphorin 7A, SEMA7A), signaling to cardiomyocytes (CPE, SLIT2, activin A), enhancement of angiogenesis (GDF6), and reduction of fibrosis (SFRP1). For several of the significantly enriched paracrine factors, for example SEMA7A, CPE, IGFBP2, and OPG, CFs have so far not been identified as a major site of production. Intriguingly, most of the secreted proteins were rather selectively expressed by the different CF populations in the heart in comparison to the other cardiac cell types (Figure 5A-B), again supporting the specificity of our experimental approach.

The numerous proteins that we found to be secreted by fibroblasts 3 and 5 days after MI clearly indicates that fibroblast activation and protein secretion is organized in a temporal fashion. This is in line with the view that fibroblasts are a dynamic cell type that achieves morphologically well-defined differentiated states following acute ischemic injury^29^. To decipher the functionality of individual factors within this secretome network, however, will not be easy and will constitute a major future task in the fibroblast field. Advanced cell culture techniques, such as organoids and in particular the heart-on-a-chip platform^42^, appear to be promising and may lead the way for the future understanding of the intercellular signaling events.

The list of infarct-induced changes in the CF secretome may hold some promise for mitigating some of the deleterious consequences of MI. Intriguingly, many of the secreted proteins identified in mice CF were confirmed at the gene expression level in mice as well as in human CF in samples of acute MI (Figure 5). Future therapeutic approaches may utilize the known anti-fibrotic activity of FMOD^43^ and SFRP1^44^ or pharmacologically neutralize the pro-fibrotic activity of BMP1, LOXL3, ECM1, and FN1 (Table 1). For some of the identified secreted proteins there is presently no cardiac function known; this includes DnaJ heat shock protein family (Hsp40) member B11 (DNAJB11), which was only detected in the miCF secretome. The secreted factors also hold promise to serve as biomarkers for CF activation in the injured heart. IGFBP2, found in the present study to be significantly elevated in the miCF secretome (Table 1), was already reported as prognostic biomarker for heart failure^45^. Similarly, OPG (*Tnfrsf11b*), a cytokine belonging to the tumor necrosis factor (TNF) receptor family which was significantly enhanced in the miCF secretome (Table 1), was also shown to be elevated during the acute phase after MI^46^ and to predict adverse cardiovascular events in stable coronary artery disease (PEACE trial)^47^. Comparing the expression level of secreted proteins between mice and human (Figure 5A-B), the following proteins may be useful plasma marker in humans for the activated fibroblast: COL1A1, COL12A1, COL16A1, FN1, LAMA2, LOX, LTBP2, PXDN, CRLF1, and SLIT2.

Communication of CF with cardiomyocytes was already suspected on the basis of single-cell analysis that identified CF as key constituent in the microenvironment to promote cardiomyocyte maturation^48^. The wide distribution of receptor expression indicates multiple potential interactions of CF secretome proteins to all cardiac cell populations (Supplemental Figure 2). The interplay between CF and monocytes has already been reported to be of pivotal importance in the diabetic heart^49^. Along this line, in co-culture experiments the secretome of cardiac stromal cells suppressed apoptosis of cardiomyocytes and preserved cardiac mitochondrial transmembrane potential^50^. Cell-to-cell communication appears also to be pathophysiologically relevant: CF from patients with idiopathic restrictive cardiomyopathy in co-culture significantly impaired relaxation velocity of healthy cardiomyocytes^51^. Now that the individual constituents of the CF secretome are identified, future studies have to decipher their specific role in intercellular cell signaling in homeostasis and disease.

In the present study we have explored the possibility to directly measure the CF secretome under in-vivo conditions by using a vector encoding ER-localized biotin ligase ER-TurboID under control of the POSTN promotor to restrict expression to activated CF (AAV9-POSTN-ER-TurboID). However, this approach showed only limited overlap with the in-vitro miCF secretome data. Firstly, POSTN, despite being specific for activated CF in the heart, is also strongly expressed in the liver^30^, so that CF-derived biotinylated proteins are “contaminated”, which we mitigated in our study by collecting the perfusate of the isolated heart. Secondly, because the number of CF within the heart is comparatively small, the intensity distribution of secreted biotinylated proteins from labeled precursor cells was only low. Thirdly, this in-vivo approach was specifically targeted to POSTN^+^ CF populations within the infarcted heart^8^, while enzymatic isolation of CF used for generation of miCF secretome data comprised all CF populations, including non-activated CFs of the remote regions. Thus, despite having shown the general feasibility of our labeling approach, future studies need to consider use of different CF promotors^52^ or use of the advantages of TurboID in a transgenic approach^53^.

In summary, this study is the first to report the secretome atlas for unstimulated (cCF) and infarct-activated fibroblasts (miCF). This secretome atlas contains aside of well-known extracellular matrix proteins numerous autocrine/paracrine proteins, of which several have previously not been assigned to CF. Most of the identified proteins are predominantly expressed in CF and a similar expression pattern was observed in human samples of AMI. Several of the identified proteins are likely to be critically involved in adverse cardiac remodeling and may serve as a rich resource for future diagnostic and therapeutic studies.

## Methods

### Animals

All experiments were performed according to the institutional and national guidelines for animal care in conformity to directive 2010/63/EU and were approved by the Landesamt für Natur-, Umwelt- und Verbraucherschutz (reference number 81-02.04.2020.A351).

Male C57Bl/6J wildtype mice were purchased from Janvier (Le Genest Isle, France) and were used at an age of 10-12 weeks and body weight of 20-25 g. Animals were housed at the Central Facility for Animal Research and Scientific Animal Welfare (ZETT) of the Heinrich Heine University Düsseldorf, Germany.

MI by ischemia/reperfusion (I/R) was induced as described previously^54^. The left anterior descending coronary artery (LAD) was ligated with an 8-0 polypropylene thread for 50 min. Ischemia was checked by ST-segment elevation on the ECG. After 50 min of ligation, the polypropylene thread was released to initiate reperfusion. For sham surgery the same procedure was used, but without LAD ligation.

### CF isolation and cultivation

CF were isolated 5 days after MI or sham surgery as described previously^17^. Mice were sacrificed by cervical dislocation and the heart was cannulated via the aorta and perfused retrogradely at 37°C with pre-warmed PBS for 3 min (constant flow of 2 ml/min) to wash out the residual blood from the coronaries. After washing, the system was switched to pre-warmed collagenase type II (Worthington Biochemical Corporation, Lakewood, USA) in PBS (1000 U/ml) for 8-9 min of digestion. To remove epicardial cells, hearts were bathed and shaken in collagenase solution which was discarded afterwards. Digested tissue was resuspended in cell culture medium (DMEM high glucose (P04-01597, PAN Biotech, Aidenbach, Germany), supplemented with 10% dialyzed FBS (Thermo Fisher Scientific, Waltham, USA) and 1% penicillin/streptomycin/glutamine (Merck Millipore, Burlington, USA)). The cell suspension was filtered using a 100 µl cell strainer and centrifuged for 1 min at 55 x*g* at 15°C to pellet cardiomyocytes. The supernatant was passed through a 40 µm cell strainer and centrifuged for 7 min at 300 *xg*. Resuspended cells were depleted for CD31^+^ and CD45^+^ cells (endothelial cells and immune cells, respectively) via magnetic depletion with Mojosort nanobeads and a Mojosort magnet (BioLegend, San Diego, USA) according to the manufacturer’s protocol and seeded in cell culture medium. On the following day, cell debris was removed by stringent washing with PBS.

### Cell viability assay

To evaluate cell viability during amino acid withdrawal and isotype labeling steps of the secretome analysis protocol, Cell Counting Kit 8 (Tebu-bio Le Perray-en-Yvelines, France), containing light-yellow tetrazolium salt that is reduced to an orange formazan by living cells, was used according to the manufacturer’s protocol. CF were isolated from healthy mouse hearts as described above and 5x10^3^ cells (in 100 µl) per well were seeded into a 96-well plate. On the next day, cells were washed with PBS and incubated in cell culture medium (see above) or depletion medium (DMEM high glucose customized without L-methionine (Met), L-arginine (Arg), and L-lysine (Lys) (PAN Biotech, Aidenbach, Germany), supplemented with 10% dialyzed FBS and 1% penicillin/streptomycin/glutamine) for up to 90 min. Subsequently, cells were washed with PBS, incubated in cell culture medium with CCK8 for 3 h, and absorbance was measured at 450 nm. To assess the effects of isotype labeling, cells were seeded as described above and incubated in depletion medium for 60 min, followed by incubation in depletion medium supplemented with 0.1 mM azidohomoalanine (AHA) (Thermo Fisher Scientific), intermediate isotypes [^13^C6] Arg (84 µg/ml) and [4,4,5,5-D4] Lys (146µg/ml), heavy isotypes [^13^C6,^15^N4] Arg (84 µg/ml) and 146µg/ml [^13^C6,^15^N2] Lys (Cambridge Isotope Laboratories, Tewksbury, USA), or combinations of AHA with intermediate and heavy isotypes for up to 24 h. CCK8 was added 3 h prior measurement. Statistical analysis was performed by Two-way ANOVA followed by Tukey’s multiple comparisons test using GraphPad Prism 9. As shown in Supplemental Figure 1, Arg/Lys/Met withdrawal up to 90 min had no effect on CF cell viability (Supplemental Figure 1A), while supplementation with AHA resulted in a significant reduction of CF cell viability after 16-24 h (Supplemental Figure 1B). Thus, incubation times of 60 min for amino acid withdrawal and 8 h for isotype labeling were chosen for the secretome analysis protocol.

### Secretome analysis in cultured CF

For secretome analysis in short-term cultured CF by liquid chromatography-tandem mass spectrometry (LC-MS/MS), a modified protocol for robust protein quantification via SILAC labeling and click chemistry according to Eichelbaum and Krijgsveld^18^ was used (see Figure 1). To reach sufficient cell numbers without expansion in cell culture, CF preparations of three mouse hearts (3 and 5 days after sham or I/R surgery, respectively) were pooled for each sample and seeded into a 10 cm cell culture dish. After cultivation in cell culture medium for 24 h at 37°C, cells were washed six times with PBS and incubated in Arg/Lys/Met depletion medium (4 ml / dish) for 1 h. Subsequently, cells were incubated in depletion medium (4 ml / dish) supplemented with 0.1 mM of the Met analog AHA and either intermediate or heavy Arg and Lys isotypes ([^13^C6] Arg (84 µg/ml), [4,4,5,5-D4] Lys (146 µg/ml) or [^13^C6, ^15^N4] Arg (84 µg/ml), [^13^C6, ^15^N2] Lys (146 µg/ml), respectively) for 8 h. Of the altogether four samples of each condition (sham / I/R surgery) at the individual time points, two were labelled with intermediate and two with heavy isotypes to exclude potential bias by the kind of labeling.

After the labeling procedure, supernatants were collected and centrifuged for 5 min at 1000 x*g* at 4°C to remove cell debris. To stabilize proteins, 1X Halt Protease-Inhibitor-Cocktail (Thermo Fisher Scientific) was added and supernatants were stored at -80°C until further use. For additional analysis of the intracellular proteome, remaining cells were washed four times with ice-cold PBS, scraped off in 1 ml PBS, and transferred to an 1.5 ml reaction tube. Cells were centrifuged for 5 min at 800 x*g* at 4°C and pellets were stored at -80°C until further use.

For enrichment of newly synthesized proteins in the supernatants, intermediate- and heavy-labelled supernatants of sham/MI sample pairs were combined and concentrated using Pierce 3 kDA protein concentrators (Thermo Fisher Scientific) at 4,000 x*g* and 4°C to a volume of about 250 µl. The Click-iT Protein Enrichment Kit (Thermo Fisher Scientific) was used according to the protocol of Eichelbaum and Krijgsveld^19^ with slight modifications. In brief, 250 µl urea buffer, 100 µl of washed agarose resin, 10 µl of dissolved Reaction Additive 2, 1 µl 10 mM Copper (II) sulfate solution, and 63 µl Reaction Additive 1 were added to the samples and samples were vortexed. To allow binding of the newly synthesized, AHA-labelled proteins to the agarose resin, samples were rotated overnight at 20 *rpm* at room temperature. On the following day, the supernatants containing non-bound proteins were removed by centrifugation for 1 min at 1000 x*g* and the resin was washed four times with 900 µl 18MΩ H2O. To reduce the resin-bound proteins, the resin was resuspended in 500 µl SDS washing buffer (warmed to 37°C) and 5 µl 1M dithiothreitol (DTT). Samples were vortexed and heated for 15 min at 70°C. After 15 min of cooling and centrifugation for 5 min at 1,000 x*g*, the resin-bound proteins were alkylated by resuspending the samples in 500 µl freshly prepared 40 mM iodoacetamide solution. After shaking the samples at 500°*rpm* for 30 min in the dark, the resin was transferred to a Mini Bio-Spin Chromatography column (Bio-Rad Laboratories, Hercules, USA), which was placed in a 2 ml reaction tube, and stringently washed by centrifugation for 30 sec at 1,000 x*g*: 10 times with 0.8 µl of SDS washing buffer, 20 times with 0.8 µl 8 M urea buffer, 20 times with 0.8 µl isopropanol, and 20 times with 0.8 µl acetonitrile.

To digest the resin-bound proteins, samples were washed twice with 500 µl digestion buffer (100 mM Tris-HCl pH 8, 10 % acetonitrile, 2 mM Ca2Cl), each time transferring the samples to a new tube. The resin was pelleted for 5 min at 1000 x*g* and supernatants was discarded, leaving about 200 µl of resin in the tube. After addition of 0.5 µg trypsin in a volume of 0.5 µl, samples were briefly vortexed and then shaken overnight at 500 *rpm* at 37°C. On the next day, the resin was pelleted for 5 min at 1000 x*g* and the peptide-containing supernatants were transferred to a new tube. The resin was resuspended in 500 µl 18MΩ H2O and again centrifuged for 5 min at 1000 x*g*. The resulting supernatants were added to the supernatants of the step before and the combined volume was filled up with 18MΩ H2O to a total volume of 1 ml to dilute the acetonitrile. To acidify the samples, 20 µl of 10% trifluoroacetic acid (TFA) were added. For desalting, Sep-Pak Cartridges (Vac 1cc (50 mg) tC18, Waters, Milford, USA) were prepared by placing them in a 20-position cartridge that was connected to a vacuum pump and washing with 900 µl acetonitrile, 300 µl of 50 % acetonitrile and 0,5 % acetic acid. After equilibrating the cartridges with 900 µl 0.1 % TFA, the samples were loaded onto the cartridges, washed with 900 µl 0.1 % TFA and 900 µl 0.5 % acetic acid, and eluted with 300 µl 50 % acetonitrile and 0.5 % acetic acid. Samples were dried using a rotational vacuum concentrator (Christ, Osterode am Harz, Germany) and stored at -80°C until further use.

Lysates of frozen CF cell pellets (see above) were prepared as described earlier^55^. Briefly, frozen CF were lysed by bead-milling in 2x 30 µl cell lysis buffer (30 mM Tris-HCl; 2 M thiourea; 7 M urea; 4% CHAPS (w/v) in water, pH 8.5). Cleared cell lysates of intermediate- and heavy-labelled CF of sham/MI sample pairs were combined (2.5 µg protein of each) and were shortly stacked into a polyacrylamide gel. Subsequently, the gel was stained with Coomassie Brilliant Blue and the protein-containing band was cut out, de-stained, reduced with dithiothreitol, alkylated with iodoacetamide and digested with trypsin overnight. Resulting peptides were extracted from the gel, dried in a vacuum concentrator and 500 ng of peptides were prepared for mass spectrometric analysis in 0.1% trifluoroacetic acid.

Peptides generated from click chemistry-enriched CF supernatant proteins and CF cell lysates, respectively, were separated on C18 material on an Ultimate3000 rapid separation LC system (Thermo Fisher Scientific) using a 2 h gradient as described earlier^55^.

Separated peptides were subsequently injected by an electrospray nano-source interface into a Fusion Lumos mass spectrometer (Thermo Fisher Scientific) operated in data-dependent, positive mode. First, survey scans were recorded in the orbitrap analyzer (resolution: 120,000, target value advanced gain control: 250,000, maximum injection time: 60 ms, scan range 200-2,000 m/z, profile mode). Second, 2-7fold charged precursor ions were selected by the quadrupole of the instrument (isolation window 1.6 m/z), fragmented by higher energy collisional dissociation (collision energy 35%), and analyzed in the linear ion-trap of the instrument (scan rate: rapid, target value advanced gain control: 10,000, maximum injection time: 50 ms, centroid mode). Cycle time was 2 s; already fragmented precursors were excluded from fragmentation for the next 60 s.

Data analysis including peptide and protein identification and MS1-based quantification was carried out with MaxQuant version 2.1.3.0 (Max Planck Institute for Biochemistry, Planegg, Germany) separately for the different sample batches with standard parameters, if not stated otherwise. MaxQuant was chosen because in our hands it shows a more reliable performance on SILAC datasets. For searches, a multiplicity of three was chosen (channel light: Arg+0 and Lys+0, channel intermediate: Arg+6, Lys+4, channel heavy: Arg+10, Lys+8) and the ‘match between runs’ function enabled. Searches were carried out on the basis of 55341 *mus musculus* proteome sequences (UP000000589) downloaded from UniProt KB on 18^th^ January 2022. Quantitative data was further processed by Perseus 1.6.6.0 (Max Planck Institute for Biochemistry, Planegg, Germany) and Excel. Here, proteins were filtered for at least two identified peptides. Intensities were normalized based on the median intensity per sample separately for the light and intermediate/heavy channels. It should be noted, that measured intensities of individual proteins can depend on available tryptic peptides, their properties, and modifications and therefore might slightly vary from the absolute protein amount. For the identification of differentially abundant proteins between CF from sham control and post-MI hearts, only proteins were considered that showed at least three valid values in one sample group. Differences were determined using the significance analysis of microarrays method^56^ based on Student’s *t*-tests and permutation-based control of the false discovery rate (FDR) which is required to account for multiple testing (FDR=5%, lysate samples: S0=0.6, secretome samples: S0=0.1). Tests were calculated on log2 transformed normalized intensity values with missing values filled in with random values drawn from a downshifted normal distribution (downshift: 1.8 standard deviations, width: 0.3 standard deviations). Identified proteins were annotated using gene ontology categories by Perseus and the secretion behavior of proteins identified were predicted by OutCyte^57^. Differences of mean values of log2 transformed normalized intensities were used for a one-dimensional annotation enrichment analysis^58^ to identify functionally related proteins which were collectively altered in their abundance.

### In-vivo-secretome analysis in POSTN^+^ CF

For in-vivo-secretome analysis of CF by LC-MS/MS, a modified protocol for cell type-selective secretome profiling according to Wei and Riley *et al.*^59^ was used (see Supplemental Figure 6). This is based on proximity labeling of proteins passing through the ER secretory pathway by ER-located biotin ligase TurboID (ER-TurboID), which is expressed under control of a cell type-specific promotor.

Plasmid pAAV-TBG-ER-TurboID was purchased from Addgene (Watertown, USA) and the TBG promotor was replaced by the POSTN promotor identified by Lindsley *et al.*^60^ to target activated CF expressing the activation marker POSTN. This pAAV-POSTN-ER-TurboID was transduced using adeno-associated virus serotype 9 (AAV9-POSTN-ER-TurboID) by injecting the virus particles (1x10^12^ genome copies) into the tail veins of mice at day 1post-MI induction by I/R surgery. Mice with I/R surgery but without virus injection were used as negative control for ER-TurboID-mediated protein labeling. Biotin was provided via the drinking water (0.5 mg/ml) to both mouse groups at days 2, 3, and 4 post-MI. At day 5 post-MI, secreted cardiac proteins were harvested by collecting the coronary effluent during Langendorff-based retrograde perfusion. To this end, explanted hearts were cannulated via the aorta and perfused with oxygenated Krebs-Henseleit buffer under constant pressure (1 m H2O). After 10 min initial perfusion to wash out residual blood, coronary effluent was collected for 60 min in 50 ml centrifuge tubes placed on ice. 1X cOmplete protease inhibitor cocktail (1 tablet / 50 ml; Roche, Basel, Switzerland) was added to the collected effluent samples to prevent protein degradation. Effluent samples (∼ 45-60 ml) were concentrated to 250 µl using Pierce 3 kDa protein concentrators (Thermo Fisher Scientific) and stored at -80°C until further use.

Enrichment of biotinylated proteins was performed according to protocols by Wei and Riley *et al.*^59^ and Cheah and Yamada^61^ using Dynabeads MyOne Streptavidin T1 (Thermo Fisher Scientific). A volume of 200 µl Dynabeads was washed twice with 1 ml lysis buffer (50 mM Tris HCl, 150 mM NaCl, 0.4% SDS, 1% NP-40, 1 mM EGTA, 1.5 mM MgCl2, 1X cOmplete protease inhibitor cocktail (1 tablet / 50 ml; Roche), pH 7.4) by centrifugation for 2 min at 2000 x*g*. The pellet was resuspended in 100 µl lysis buffer and 200 µl of the concentrated effluent was added. Samples were rotated overnight at 20 *rpm* and 4°C. Samples were washed with 1 ml lysis buffer, 1 ml washing buffer (50 mM Tris HCl, 2% SDS, pH 7.5) and twice with 1 ml lysis buffer. For elution, samples were incubated for 5 min at 95°C after addition of 15 µl of a 25 mM biotin solution. Samples were centrifuged for 2 min at 2000 x*g* and supernatants containing biotinylated proteins were transferred into 2 ml reaction tubes and stored at -80°C until further use.

For LC-MS/MS analysis, eluted proteins were shortly stacked into a polyacrylamide gel. After staining with Coomassie Brilliant Blue, the protein-containing bands were cut out, de-stained, reduced with dithiothreitol, alkylated with iodoacetamide and digested with trypsin overnight as described^55^. After extraction from the gel, the peptides were dried in a vacuum concentrator and 1/3rd of peptides were prepared for mass spectrometric analysis in 0.1% trifluoroacetic acid.

Samples were analyzed on a Fusion Lumos mass spectrometer (Thermo Fisher Scientific) in data-dependent, positive mode, in a similar setting as described above for secreted proteins but with some modifications: the gradient length for peptide separation was only 1 h and some settings for spectrum acquisition were different. Full scans were recorded in the orbitrap (resolution: 120,000, scan range: 200-2000 m/z, maximum injection time: 60 ms, automatic gain control target: 400,000, profile mode), 2-7fold charged precursors selected for quadrupole-based isolation (1.6 m/z isolation window), fragmented by higher-energy collisional dissociation (collision energy 35%) and fragment spectra recorded in the linear ion trap of the instrument (scan rate: rapid, scan range: auto, maximum injection time: 150 ms, automatic gain control target: 10,000, centroid mode). Cycle time was 2 s and already isolated precursors were excluded from isolation for the next 60 s.

Data analysis was carried out with Proteome Discoverer 2.4.1.15 (Thermo Fisher Scientific) in order to profit from an enhanced sensitivity in comparison to MaxQuant-based searches. For protein identification, sequences as indicated above for cultured CF were used for protein identification by Sequest HT (10 ppm and 0.6 Da mass deviation for precursor and fragment masses, respectively; variable modification: methionine oxidation, N-terminal acetylation and methionine loss; fixed modification: carbamidomethyl on cysteines). Proteins and peptides were accepted based on the Percolator node with a false discovery rate of 1%. The precursor quantification node was used for peptide and protein quantification. Only “high confidence” proteins were reported, which were identified with at least two different peptides and did not occur in the list of potential contaminant sequences. Statistical data analysis was carried out as described above for isolated CF samples, using an S0 of 0.6 and 5% FDR on log2 transformed normalized intensities. To further increase confidence, proteins detected in <4 of the 6 samples in the virus-injected mouse group were excluded from further analysis.

### Immunofluorescence analysis

To analyze the distribution of ER-TurboID expression in the healthy and post-MI heart, AAV9-POSTN-ER-TurboID virus particles were injected into the tail vein (1x10^12^ genome copies) 2 days after I/R or sham surgery. Biotin was provided via the drinking water at days 4, 5, and 6 post-MI. At day 7 post-MI, mice were sacrificed and hearts were embedded in tissue freezing medium. Cryostat sections were prepared and were fixed with 4% paraformaldehyde (PFA) for 15 min. Immunostaining was performed using primary antibodies specific for POSTN (1:100; OriGene, Rockville, USA) and V5 tag (1:100, Thermo Fisher Scientific) to detect V5-tagged ER-TurboID, and secondary antibodies AF488-coupled goat-anti-rabbit IgG and AF594-coupled goat-anti-mouse IgG (1:1,000, Thermo Fisher Scientific) in presence of 0.2% saponin. Cover slips were mounted with ProLong Gold Antifade Reagent with DAPI to label cell nuclei. Fluorescence microscopy was performed using a BX61 fluorescence microscope (Olympus, Tokyo, Japan). Images were processed for publication using ImageJ/Fiji^62^.

### Metabolic RNA sequencing

To quantify newly synthesized and existing RNA transcripts in CF, thiol(SH)-linked alkylation for the metabolic sequencing (SLAMseq) using the SLAMseq Explorer and Kinetics Kit (Lexogen, Vienna, Austria) was performed. CF isolated from healthy hearts were seeded into a 24-well plate (30,000 cells / well, two wells / sample) and grown until confluency. Then, cells were cultured in culture medium with the uridine analog 4-Thiouridine (S4U, 0,1 mM) to allow S4U incorporation into newly synthesized RNA transcripts. As negative control, cells were incubated in parallel without addition of S4U. After 12 h, CF were lysed by TRIsure (Meridian Bioscience, Cincinnati, USA). RNA isolation and alkylation by iodoacetamide, which results in incorporation of a guanine instead of an adenine at S4U nucleotides during downstream NGS library preparation, were performed according to the kit manufacturer’s protocol including the optional addition of unique molecular identifiers (UMI) for later deduplication.

The QuantSeq 3’ mRNA-Seq Library Prep Kit FWD (Lexogen, Vienna, Austria) was used to prepare NGS libraries with the optional UMI added for duplicate removal in later stages of the analysis. At least 500 pg of alkylated RNA samples were used as input. High-throughput sequencing of the libraries was performed using the Illumina NextSeq1000/2000 system (Illumina Inc., San Diego, USA) with a target of 50mio single-end, 75bp long-reads per sample. Demultiplexing was performed using bcl2fastq2 as part of a snakemake-based pipeline (https://github.com/WestGermanGenomeCenter/bcl2fastq2_Pipeline). Sequence quality was checked using FastQC, MultiQC and the SLAM-DUNK software^63^ utilizing its utrrates function. T>C conversions, which result from the incorporation of guanine at alkylated S4U nucleotides during NGS library preparation, were detected and quantified with one minor addition. Briefly, the previously added UMI were moved from the reads sequence into the read header using umi-tools (https://umi-tools.readthedocs.io/en/latest/) extract function, discarding all reads without complete UMI and TATA-spacer sequence before mapping. After the mapping step of SLAM-DUNK (using mm39 as the reference genome), mapped reads were deduplicated with umi-tools dedup. Further analysis of the SLAM-DUNK output data was performed using R. Transcripts without T detection in a sample were removed. A beta-binomial test was performed according to manufacturer’s instructions using the bb.test function. *P* values were calculated to detect transcripts with significantly increased T>C conversion rate (*P* < 0.05).

### Single-cell / single-nucleus RNA sequencing data analysis

Single-cell RNA sequencing (scRNAseq) and single-nucleus RNA sequencing (snRNAseq) data were processed using the R Seurat package^64^ (v3.0).

For comparison of gene expression among cardiac cell types including cardiomyocytes, an snRNAseq data set of cardiac cells isolated from an healthy heart of a 12-week-old mouse^22^ (ArrayExpress E-MTAB-7869, sample Y1) was used.

For comparison of gene expression in CF 5 days after sham and MI surgery or among the cardiac cell types 5 days post-MI, we re-analyzed scRNAseq data sets previously published by us^28^. Names of CF populations were annotated according to expression of CF population markers on basis of Farbehi *et al.*^65^ and Shi *et al.*^66^.

To assess gene expression in human healthy hearts, post-MI hearts and hearts with cardiomyopathy, we analyzed snRNAseq and cellular indexing of transcriptomes and epitopes by sequencing (CITE-seq) data sets recently generated^23–25^.

### Statistics and Reproducibility

The methods used are described in detail in the respective ‘Secretome analysis in cultured CF’, ‘In-vivo-secretome analysis in POSTN^+^ CF’, and ‘Metabolic RNA sequencing’ sections and in the figure legends (where appropriate). *P* values of <0.05 were considered statistically significant. For secretome analysis in cultured CF, experiments were performed with biological replicates in terms of CF preparations from different animals. For in-vivo-secretome analysis, experiments were performed with biological replicates in terms of coronary effluents collected from different animals. Numbers of biological replicates are stated in the figure legends where appropriate.

## Data availability

Mass spectrometry data have been deposited to the ProteomeXchange Consortium via the PRIDE^67^ partner repository with the dataset identifiers PXD046238 and PXD053791.

## Acknowledgements

This work was funded by the Deutsche Forschungsgemeinschaft (DFG, German Research Foundation) *–* 458365199. This work was supported by the DFG Research Infrastructure West German Genome Center, project 407493903, as part of the Next Generation Sequencing Competence Network, project 423957469.

## Author contributions

J.B.: Conceptualization, methodology, validation, formal analysis, investigation, writing – original draft, writing – review & editing, visualization, project administration. G.P.: methodology, validation, formal analysis, investigation, resources, data curation, writing – original draft, writing – review & editing, visualization, project administration. A.J..: Methodology, Resources. M.B.: Methodology, Resources. Z.D.: Investigation, resources. R.Z.: Investigation. J.St.: Investigation. T.W.: Methodology, formal analysis, investigation. D.R.: Methodology, formal analysis, investigation, data curation. T.L.: Formal analysis, investigation, resources, data curation. C.A.: Investigation, resources. J.A.A.: Formal analysis, resources, data curation, visualization. K.J.L.: Resources, supervision. K.K.: Methodology, resources, supervision. P.M.: Methodology, resources, supervision. K.S.: Methodology, resources, supervision, writing – original draft. J.H.: Conceptualization, formal analysis, investigation, writing – original draft, writing – review & editing, visualization, project administration, funding acquisition. J.Sch.: Conceptualization, supervision, writing – original draft, writing – review & editing, project administration, funding acquisition.

## Competing interests

The authors declare no competing interests.

## Notes

### Competing Interest Statement

The authors have declared no competing interest.

