## Supplemental Figures and Tables for "Defining the cardiac fibroblast secretome in the healthy and infarcted mouse heart"

### Supplemental Material

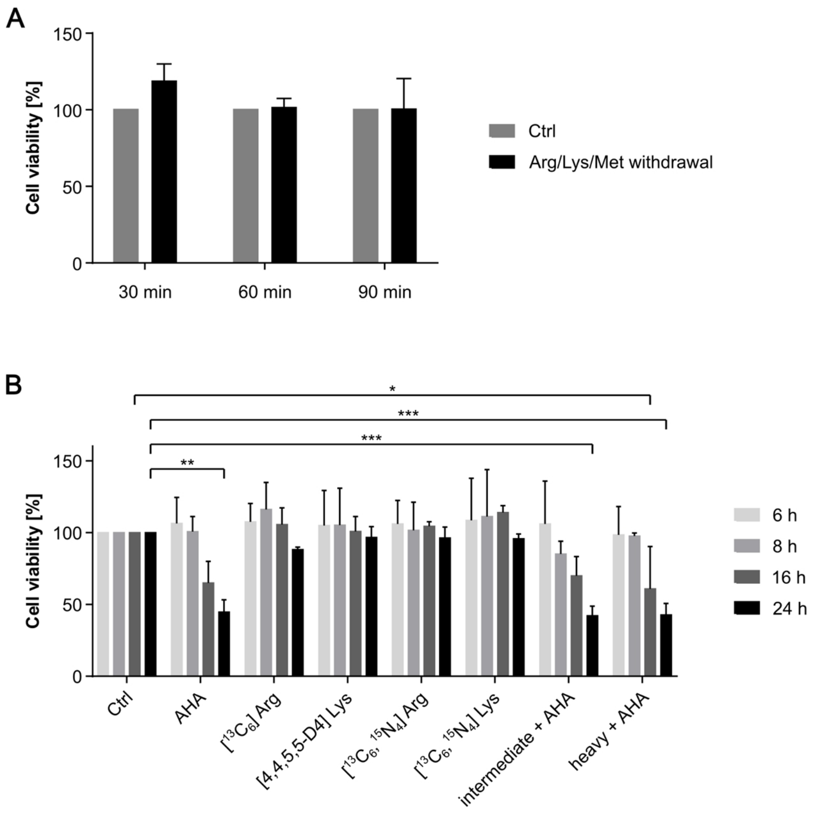


**Supplemental Figure 1: Effects of AHA and Arg/Lys isotype labeling on CF cell viability.**
**A)** CF isolated from healthy mouse hearts were incubated in conventional cell culture medium (Ctrl) and depletion medium without L-methionine (Met), L-arginine (Arg), and L-lysine (Lys) (Arg/Lys/Met withdrawal). **B)** After incubation in depletion medium without Met, Arg, and Lys for 1 h, CF were incubated in depletion medium supplemented with AHA, intermediate isotypes [$^{13}C_6$] Arg and [4,4,5,5-$D_4$] Lys, heavy isotypes [$^{13}C_6$,$^{15}N_4$] Arg and [$^{13}C_6$,$^{15}N_2$] Lys, or combinations of AHA with intermediate and heavy isotypes for the indicated time points. As control (Ctrl), CF were incubated in conventional cell culture medium. All media contained 10% FBS. At the indicated time points, CF cell viability was quantified via Cell Counting Kit-8. The viability of Ctrl cells was set to 100%. Data are shown as mean ± SEM (n=3). Two-way ANOVA followed by Tukey's multiple comparisons test, *$P$<0.05, **$P$<0.01, ***$P$<0.001.

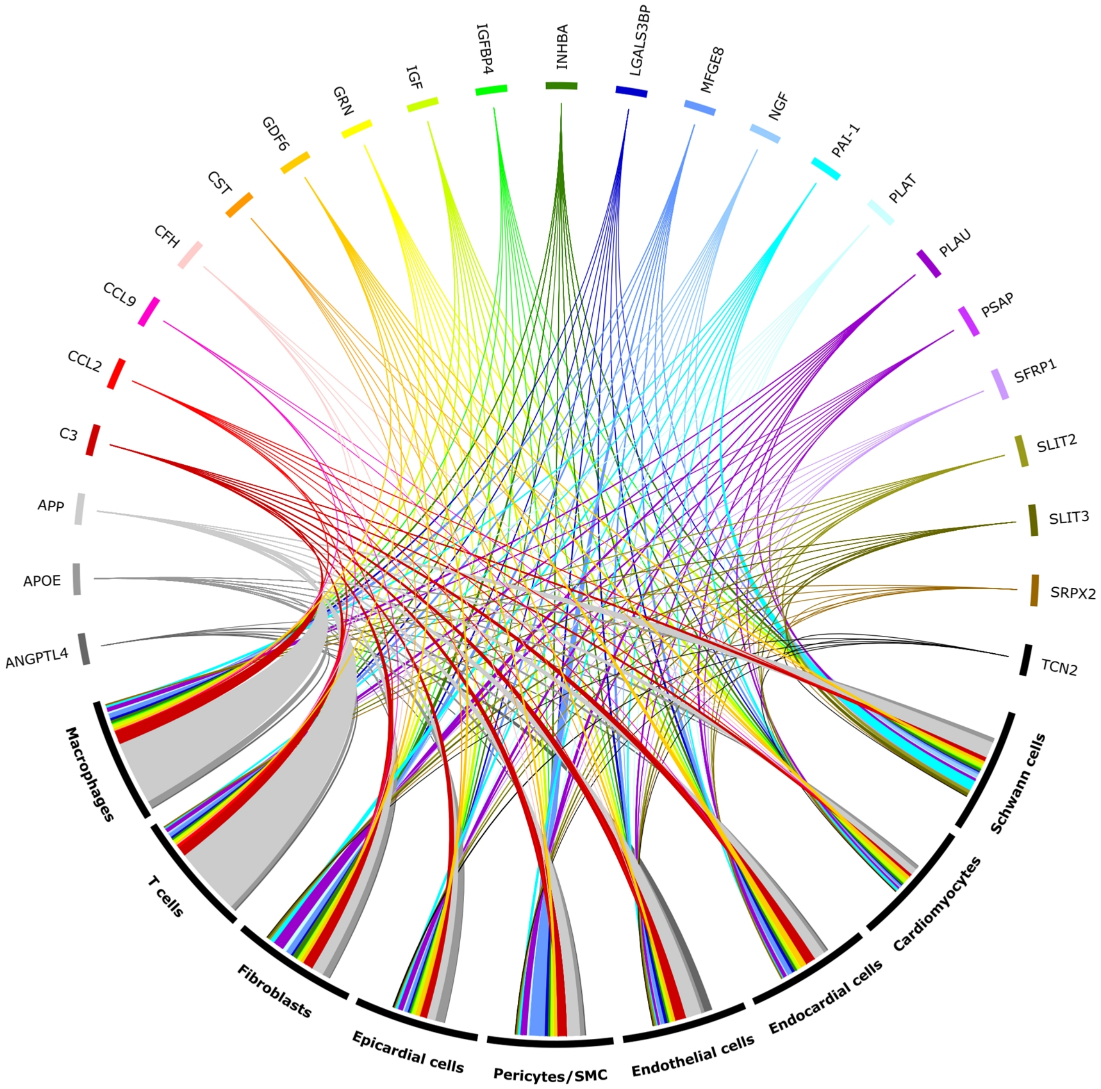


**Supplemental Figure 2: Potential cell-cell-communication mediated by cCF paracrine/autocrine factors.** Receptors of cCF paracrine/autocrine factors were selected from the ligand-receptor database of CellTalkDB[1] and snRNAseq data from n=1 healthy mouse heart published by Vidal *et al.*[2] (sample Y1, 3,790 cells) were analyzed for their expression. The Circos[3] plot visualizes potential interactions between cCF paracrine factors and their receptor(s) in the different cardiac cell types. The thickness of the lines indicates the receptor expression levels in the respective cell types. Cell types are ordered according to population size.

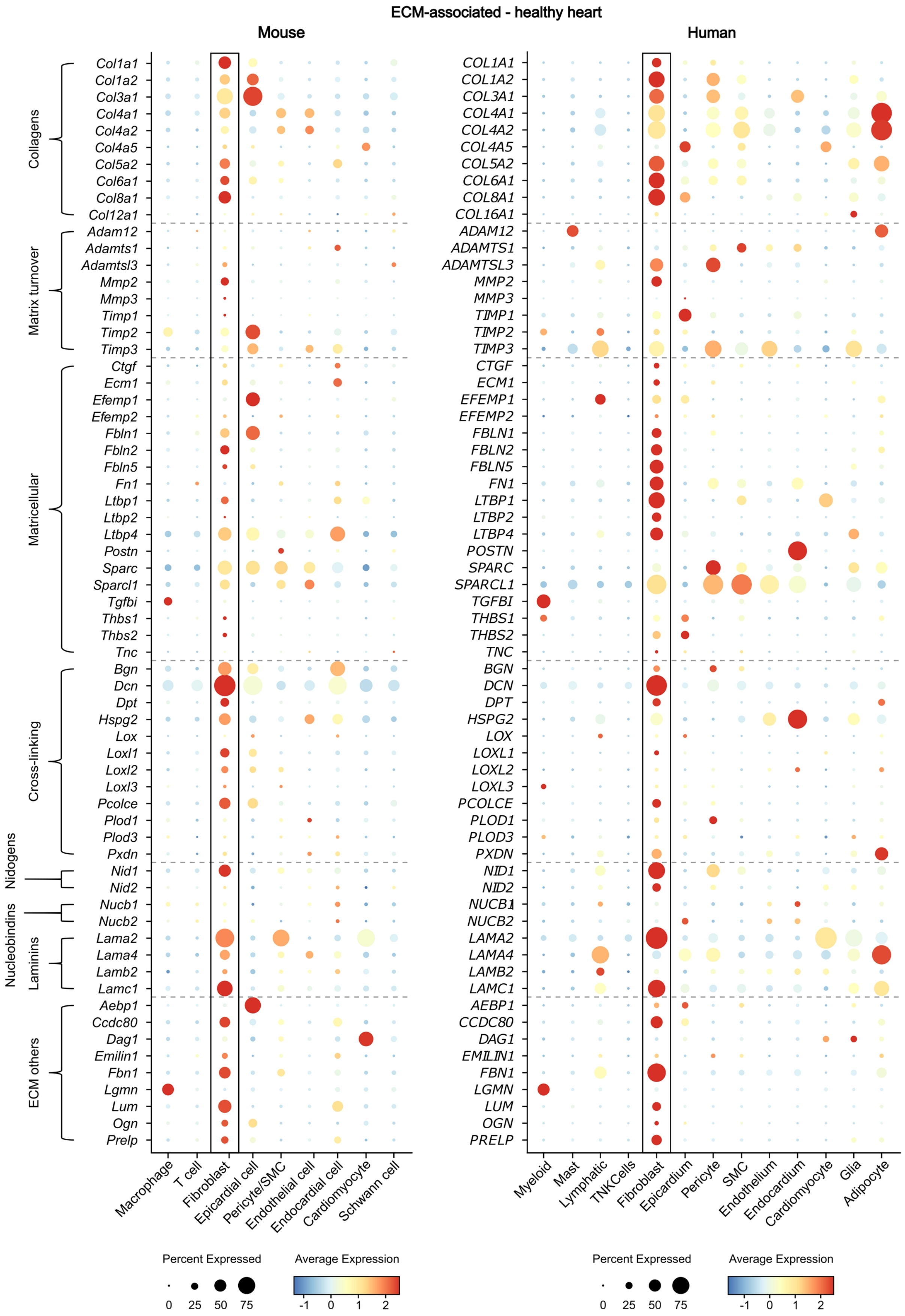

ECM-associated - healthy heart
Mouse
Human
Collagens
Matrix turnover
Matricellular
Cross-linking
Nidogens
Nucleobindins
Laminins
ECM others
Col1a1
Col1a2
Col3a1
Col4a1
Col4a2
Col4a5
Col5a2
Col6a1
Col8a1
Col12a1
Adam12
Adamts1
Adamtsl3
Mmp2
Mmp3
Timp1
Timp2
Timp3
Ctgf
Ecm1
Efemp1
Efemp2
Fbln1
Fbln2
Fbln5
Fn1
Ltbp1
Ltbp2
Ltbp4
Postn
Sparc
Sparcl1
Tgfbi
Thbs1
Thbs2
Tnc
Bgn
Dcn
Dpt
Hspg2
Lox
Loxl1
Loxl2
Loxl3
Pcolce
Plod1
Plod3
Pxdn
Nid1
Nid2
Nucb1
Nucb2
Lama2
Lama4
Lamb2
Lamc1
Aebp1
Ccdc80
Dag1
Emilin1
Fbn1
Lgmn
Lum
Ogn
Prelp
Macrophage
T cell
Fibroblast
Epicardial cell
Pericyte/SMC
Endothelial cell
Endocardial cell
Cardiomyocyte
Schwann cell
COL1A1
COL1A2
COL3A1
COL4A1
COL4A2
COL4A5
COL5A2
COL6A1
COL8A1
COL16A1
ADAM12
ADAMTS1
ADAMTSL3
MMP2
MMP3
TIMP1
TIMP2
TIMP3
CTGF
ECM1
EFEMP1
EFEMP2
FBLN1
FBLN2
FBLN5
FN1
LTBP1
LTBP2
LTBP4
POSTN
SPARC
SPARCL1
TGFBI
THBS1
THBS2
TNC
BGN
DCN
DPT
HSPG2
LOX
LOXL1
LOXL2
LOXL3
PCOLCE
PLOD1
PLOD3
PXDN
NID1
NID2
NUCB1
NUCB2
LAMA2
LAMA4
LAMB2
LAMC1
AEBP1
CCDC80
DAG1
EMILIN1
FBN1
LGMN
LUM
OGN
PRELP
Myeloid
Mast
Lymphatic
TNKCells
Fibroblast
Epicardium
Pericyte
SMC
Endothelium
Endocardium
Cardiomyocyte
Glia
Adipocyte
Percent Expressed
0
25
50
75
Average Expression
-1
0
1
2
Percent Expressed
0
25
50
75
Average Expression
-1
0
1
2

**Supplemental Figure 3: Gene expression of ECM-associated proteins identified in secretome analysis of cCF within the major cardiac cell populations as assessed by single-cell transcriptomics.**

SnRNAseq data of n=1 healthy mouse heart published by Vidal *et al.*[2] (sample Y1, 3,790 cells; left panel) and snRNAseq data from n=25 human hearts of healthy donors[4–6] (right panel) were re-analyzed. Cellular distribution of gene expression of ECM-associated cCF secretome proteins is visualized as dot plot.

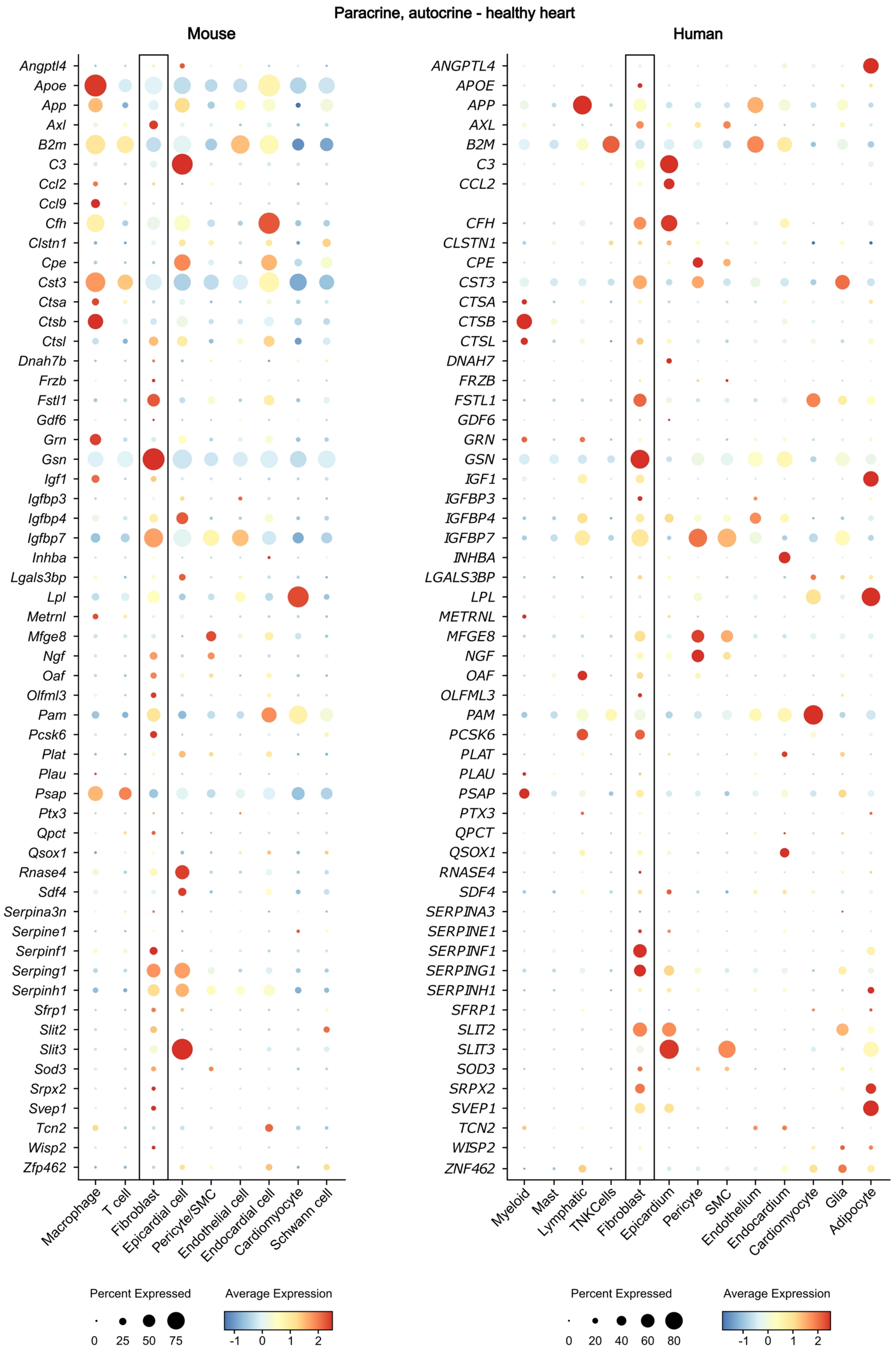

Paracrine, autocrine - healthy heart
Mouse
Human
Angptl4
Apoe
App
Axl
B2m
C3
Ccl2
Ccl9
Cfh
Clstn1
Cpe
Cst3
Ctsa
Ctsb
Ctsl
Dnah7b
Frzb
Fstl1
Gdf6
Grn
Gsn
Igf1
Igfbp3
Igfbp4
Igfbp7
Inhba
Lgals3bp
Lpl
Metrnl
Mfge8
Ngf
Oaf
Olfml3
Pam
Pcsk6
Plat
Plau
Psap
Ptx3
Qpct
Qsox1
Rnase4
Sdf4
Serpina3n
Serpine1
Serpinf1
Serping1
Serpinh1
Sfrp1
Slit2
Slit3
Sod3
Srpx2
Svep1
Tcn2
Wisp2
Zfp462
Macrophage
T cell
Fibroblast
Epicardial cell
Pericyte/SMC
Endothelial cell
Endocardial cell
Cardiomyocyte
Schwann cell
ANGPTL4
APOE
APP
AXL
B2M
C3
CCL2
CFH
CLSTN1
CPE
CST3
CTSA
CTSB
CTSL
DNAH7
FRZB
FSTL1
GDF6
GRN
GSN
IGF1
IGFBP3
IGFBP4
IGFBP7
INHBA
LGALS3BP
LPL
METRNL
MFGE8
NGF
OAF
OLFML3
PAM
PCSK6
PLAT
PLAU
PSAP
PTX3
QPCT
QSOX1
RNASE4
SDF4
SERPINA3
SERPINE1
SERPINF1
SERPING1
SERPINH1
SFRP1
SLIT2
SLIT3
SOD3
SRPX2
SVEP1
TCN2
WISP2
ZNF462
Myeloid
Mast
Lymphatic
TNKCells
Fibroblast
Epicardium
Pericyte
SMC
Endothelium
Endocardium
Cardiomyocyte
Glia
Adipocyte
Percent Expressed
0 25 50 75
Average Expression
-1 0 1 2
Percent Expressed
0 20 40 60 80
Average Expression
-1 0 1 2

**Supplemental Figure 4: Gene expression of paracrine/autocrine factors identified in secretome analysis of cCF within the major cardiac cell populations as assessed by single-cell transcriptomics.**
SnRNAseq data of n=1 healthy mouse heart published by Vidal et al.[2] (sample Y1, 3,790 cells; left panel) and snRNAseq data from n=25 human hearts of healthy donors[4–6] (right panel) were re-analyzed. Cellular distribution of gene expression of paracrine/autocrine cCF secretome proteins is visualized as dot plot.

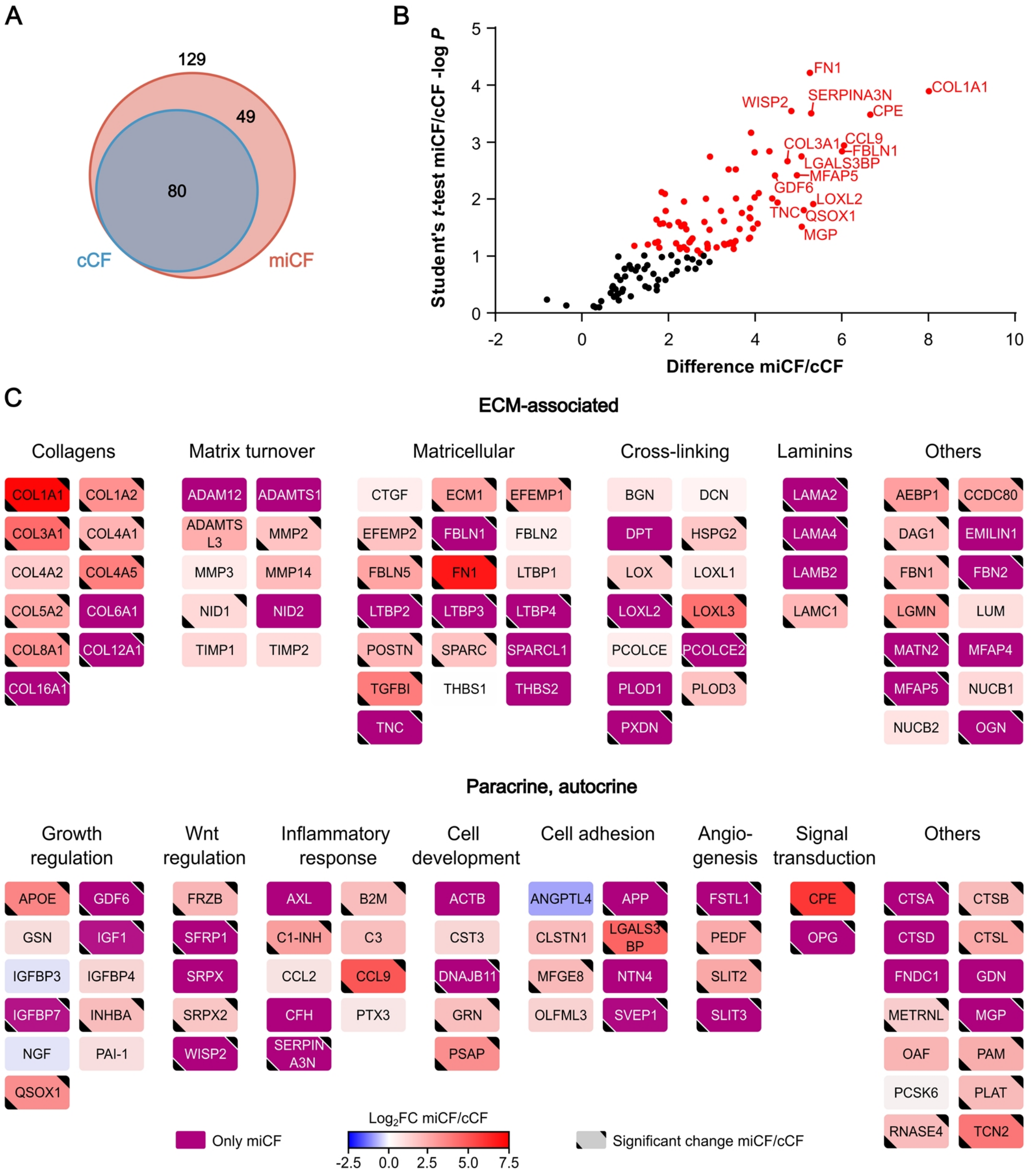


**Supplemental Figure 5: MI-induced changes of protein intensities 3 days after infarction.**
LC-MS/MS analysis identified 129 secreted proteins from post-MI CF (miCF) isolated from mouse hearts 3 days after I/R surgery (n=4; source data in Supplemental Data 1). These data were compared to the basal secretome data (80 proteins) of cCF isolated from sham-operated hearts 3 days after surgery (n=4; source data in Supplemental Data 1). **A)** Venn diagram[7] of identified proteins. **B)** Volcano plot of differentially secreted proteins. Proteins with significantly different intensities in cCF and miCF samples (Student's *t*-test-based SAM analysis, 5% FDR, $S_0$=0.1) are highlighted in red (79 proteins). For statistical significance analysis of secreted proteins which were only detected in miCF, an imputation approach of missing base values was performed, using values taken from a downshifted normal distribution (details in Methods). Names of top 15 proteins with highest difference between cCF and miCF are annotated. The shown difference refers to the difference of group mean values of $\log_2$ transformed intensities. **C)** Log2 fold changes (FC) of protein intensities between miCF and cCF samples visualized with Cytoscape[8]. Proteins were grouped in subcategories ECM-associated proteins and paracrine/autocrine factors.

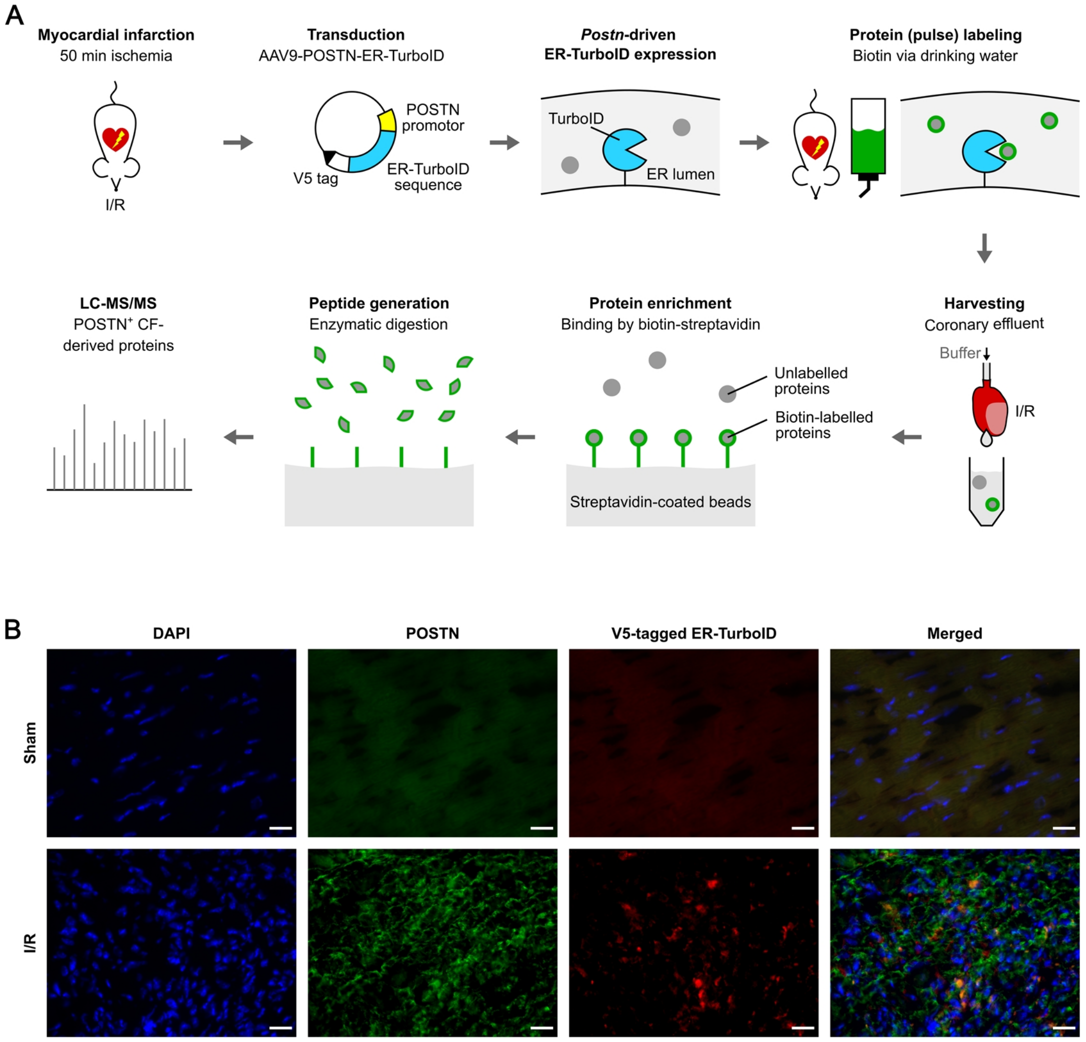


**Supplemental Figure 6: Workflow of in-vivo-secretome analysis of POSTN⁺ CF.**
**A)** At day 1 post MI (50 min ischemia/reperfusion, IR), adeno-associated virus serotype 9 (AAV9) carrying an expression vector encoding V5-tagged ER-localized biotin ligase TurboID (ER-TurboID) under control of the POSTN promotor (AAV9-POSTN-ER-TurboID) was injected i.v.. To allow TurboID expressed in MI-activated POSTN⁺ CF to biotinylate proximate proteins passing through the ER secretory pathway, biotin was provided via the drinking water for 3 consecutive days before protein harvest. At 5 days post-MI, secreted cardiac proteins were collected in the coronary effluent during Langendorff-based retrograde perfusion. POSTN⁺ CF-derived, biotin-labelled proteins were bound to streptavidin-coated beads. After stringent washing to remove non-bound proteins, remaining proteins were enzymatically digested and eluted peptides were applied to LC-MS/MS. **B)** Expression of POSTN and V5-tagged ER-TurboID as assessed by immunofluorescence analysis. AAV9-POSTN-ER-TurboID was injected 2 days after MI or sham surgery (n=1 each). Biotin was provided for 3 consecutive days before sacrificing the mice at day 7 post-MI and preparing cryosections. Immunofluorescence analysis was performed with antibodies specific for POSTN and the V5-tag of ER-TurboID. DAPI was used to label nuclei. Representative images of the infarct border zone are shown.

**Supplemental Table 1: Reported functions of paracrine/autocrine factors secreted from cCF**

Proteins with known functions that were found in the secretome of non-activated CF (cCF) isolated from the mouse hearts 5 days after sham surgery were selected by PubMed search. Proteins are listed in decreasing secretome protein intensity. Quantitative information of the listed proteins can be found in Figure 2 and Supplemental Data 1.

| Protein; *gene* | Function | Ref. |
|---|---|---|
| **Plasminogen activator inhibitor 1 (PAI-1); *Serpine1*** | Central function in thrombo-inflammation | 9 |
| **Gelsolin (GSN); *Gsn*** | Important mediator of cardiac fibrosis | 10 |
| **Insulin like growth factor binding protein 4 (IGFBP4); *Igfbp4*** | Promotes cardiogenesis | 11 |
| **Pentraxin-related protein 3 (PTX3); *Ptx3*** | Essential component of the humoral arm of the innate immune system | 12 |
| **Complement C3 (C3); *C3*** | Important component of complement system | 13 |
| **Chemokine (C-C motif) ligand 2 (CCL2); *Ccl2*** | Regulates migration and infiltration of a wide range of immune cells | 14 |
| **Pigment epithelium-derived factor (PEDF); *Serpinf1*** | Potent inhibitor of angiogenesis | 15 |
| **Beta-nerve growth factor (NGF); *Ngf*** | Beneficial actions on cardiomyocytes; angiogenesis | 16 |
| **Angiopoietin-like 4 (ANGPTL4); *Angptl4*** | Antiangiogenic modulatory factor | 17 |
| **Proprotein convertase subtilis/kexin 6 (PCSK6); *Pcsk6*** | Modulates cardiomyocyte senescence | 18 |
| **Follistatin-related protein 1 (FSTL1); *Fstl1*** | Critical for homeostasis of vascular wall | 19 |
| **Superoxide dismutase (SOD3); *Sod3*** | Redox signaling; modulation of inflammatory response | 20 |
| **Tyrosine-protein kinase receptor UFO (AXL); *Axl*** | Mediates inflammation | 21 |

**Supplemental Table 2: Intracellular protein and mRNA transcript levels of proteins of the cCF secretome.** The cCF proteome in cell lysates harvested together with the supernatants for the secretome analysis after 8 h of SILAC labeling was assessed by LC-MS/MS (n=4; source data in Supplemental Data 2). Bulk transcriptome analysis with detection of newly synthesized transcripts (thiol-linked alkylation for metabolic sequencing, SLAMseq) was performed in CF isolated from healthy mouse hearts (n=3; source data in Supplemental Data 3). For transcript labeling, CF were incubated for 12 h with 4-thiouridine (S4U). In subsequent sample processing, incorporated S4U in newly synthesized transcripts resulted in T>C conversions that were quantified by sequencing. For selected proteins of the cCF secretome, their protein levels (intensities with SILAC label, Int) in the secretome and proteome, as well as the mRNA transcript levels (counts per million, CPM) and transcript T>C conversion rates are shown. Data are reported as mean ± SD. n.d., not detected.

| **Protein/*gene*** | **Secretome [Int]** | | **Proteome [Int]** | | **Transcriptome [CPM]** | | **Transcriptome [T>C conversion rate]** | |
|---|---|---|---|---|---|---|---|---|
| **PAI1/*Serpine1*** | **7.06E+09** | ±2.37E+09 | **4.75E+06** | ±3.11E+06 | **340.57** | ±180.03 | **0.050** | ±0.001 |
| **GSN/*Gsn*** | **9.31E+08** | ±2.49E+08 | **n.d.** | | **42.81** | ±33.62 | **0.015** | ±0.003 |
| **IGFBP4/*Igfbp4*** | **3,81E+08** | ±5.13E+07 | **n.d.** | | **150.94** | ±58.19 | **0.014** | ±0.002 |
| **PTX3/*Ptx3*** | **3.51E+08** | ±1.04E+08 | **3.24E+07** | ±2.18E+07 | **60.26** | ±49.01 | **0.045** | ±0.003 |
| **C3/*C3*** | **1.34E+08** | ±6.82E+07 | **n.d.** | | **51.45** | ±25.28 | **0.007** | ±0.002 |
| **CCL2/*Ccl2*** | **8.71E+07** | ±3.45E+07 | **n.d.** | | **36.95** | ±33.56 | **0.052** | ±0.003 |
| **NGF/*Ngf*** | **2.11E+07** | ±5.77E+06 | **n.d.** | | **9.58** | ±8.36 | **n.d.** | |
| **CTSB/*Ctsb*** | **1.83E+07** | ±2.42E+07 | **2.10E+07** | ±4.50E+06 | **76.83** | ±38.53 | **0.011** | ±0.003 |
| **CCL9/*Ccl9*** | **1.82E+07** | ±1.21E+07 | **n.d.** | | **5.83** | ±4.31 | **0.028** | ±0.006 |
| **IGF1/*Igf1*** | **1.57E+07** | ±4.92E+06 | **n.d.** | | **53.84** | ±40.86 | **0.039** | ±0.002 |
| **ANGPTL4/*Angptl4*** | **1.50E+07** | ±7.18E+06 | **n.d.** | | **3.912** | ±2.35 | **0.031** | ±0.004 |
| **CFH/*Cfh*** | **1.35E+07** | ±1.85E+07 | **n.d.** | | **80.87** | ±63.58 | **0.015** | ±0.000 |
| **PCSK6/*Pcsk6*** | **1.30E+07** | ±5.56E+06 | **n.d.** | | **19.82** | ±12.56 | **0.019** | ±0.005 |
| **FSTL1/*Fstl1*** | **9.91E+06** | ±3.45E+06 | **n.d.** | | **246.95** | ±89.34 | **0.030** | ±0.000 |
| **LPL/*Lpl*** | **3.37E+06** | ±5.61E+05 | **n.d.** | | **148.99** | ±85.05 | **0.029** | ±0.003 |
| **AXL/*Axl*** | **2.64E+06** | ±1.44E+06 | **n.d.** | | **21.12** | ±10.84 | **0.040** | ±0.004 |
| **HSP47/*Serpinh1*** | **1.84E+06** | ±1.12E+06 | **1.63E +09** | ±5.45E+08 | **235.86** | ±117.12 | **0.022** | ±0.002 |

**Supplemental Table 3: Secretome proteins of miCF at day 3 and day 5 post-MI that were significantly changed in comparison to cCF from sham-operated hearts.**

Secretome data obtained from miCF isolated 3 days after MI and secretome data obtained from miCF isolated 5 days after MI were matched in terms of significantly upregulated proteins compared to respective sham controls (n=4 each, source data in Supplemental Data 1).

| Proteins significantly upregulated | | | | |
|---|---|---|---|---|
| **only at day 3 post-MI** | | | **both at day 3 and 5 post-MI** | **only at day 5 post-MI** |
| AEBP1 | FBLN5 | NID1 | COL1A1 | ACTG1 |
| APOE | FBN1 | OGN | COL12A1 | BMP1 |
| APP | FBN2 | PAM | COL16A1 | CRLF1 |
| B2M | FRZB | PCOLCE2 | CPE | FMOD |
| C1-INH | FSTL1 | PEDF | DNAJB11 | IGFBP2 |
| CCDC80 | GRN | PLAT | ECM1 | PLOD1 |
| CCL9 | HSPG2 | PLOD3 | FN1 | SEMA7A |
| COL1A2 | IGF1 | PSAP | GDF6 | TIMP3 |
| COL3A1 | IGFBP7 | QSOX1 | INHBA | |
| COL4A1 | LAMC1 | RNASE4 | LAMA2 | |
| COL4A5 | LGALS3BP | SERPINA3N | LAMA4 | |
| COL5A2 | LGMN | SLIT3 | LOX | |
| COL8A1 | LOXL2 | SPARC | LOXL3 | |
| CTSA | LTBP3 | SRPX2 | LTBP2 | |
| CTSB | LTBP4 | SVEP1 | MATN2 | |
| CTSL | METRNL | TCN2 | POSTN | |
| DAG1 | MFAP5 | TGFBI | PXDN | |
| EFEMP1 | MFGE8 | TNC | SFRP1 | |
| EFEMP2 | MGP | WISP2 | SLIT2 | |
| FBLN1 | MMP2 | | OPG | |

**Supplemental Table 4: Secretome proteins of cultured miCF and POSTN$^+$ CF in-vivo 5 days post-MI.** Identified proteins in the secretome of miCF isolated 5 days after MI (n=4; source data in Supplemental Data 1) and in the in-vivo-secretome of POSTN$^+$ CF collected in the coronary effluent at day 5 post-MI (n=6; source data in Supplemental Data 4) were matched. Proteins that were significantly enriched in the effluent of AAV9-POSTN-ER-TurboID-transduced mice in comparison to non-transduced control mice 5 days post-MI (n=6; source data in Supplemental Data 4) are marked with an asterisk.

| Identified proteins detected | | | | | | |
|---|---|---|---|---|---|---|
| only in cultured miCF | | | both in cultured miCF and coronary effluent | only in coronary effluent | | |
| ACTG1 | EFEMP2 | OPG | APOE* | A1AT2* | EGFR* | ITIH2 |
| ADAM12 | EMILIN1 | PAI-1 | BGN | A1AT3* | EIF4A1* | ITIH3 |
| ADAM15 | FBN1 | PAM | C1-INH* | A1AT4* | EPHX2 | ITIH4* |
| ADAMTS1 | FMOD | PCOLCE2 | C3* | A1AT5* | EZR | KLKB1* |
| ADAMTS2 | FNDC1 | PCSK6 | CFH* | A2AP* | F11* | KNG1* |
| ADAMTSL3 | FRZB | PLAT | COL3A1 | ACTA2 | F12* | KNG2* |
| ADM | FSTL1 | PLAU | CP* | ACTB | F13B* | KRT36 |
| AEBP1 | GAS6 | PLOD1 | DCN | ACTN2 | F2* | KRT76 |
| ANGPTL4 | GDF6 | PLOD3 | FBLN1 | AFM* | FABP3 | LDHA |
| APP | GDN | PRELP | FBLN2* | AGT* | FABP4 | LDHB |
| ASPN | HSP47 | PROS1 | FBLN5 | AI182371* | FETUB* | LIFR* |
| AXL | IGF1 | PSAP | FN1 | ALB* | FGA | LRG1* |
| B2M | IGFBP2 | PTX3 | GRN* | ALDOA | FGB | LYZ1 |
| BMP1 | IGFBP3 | QPCT | GSN | AMBP* | FGG | LYZ2 |
| CCDC80 | IGFBP4 | RNASE4 | HSPG2 | ANT3 | FHL2 | MB |
| CCL2 | IGFBP7 | SDF4 | LAMA4 | ANXA2 | GAPDH | MBL1* |
| CCL9 | INHBA | SEMA7A | LAMB1 | APOA1* | GC* | MBL2* |
| CLSTN1 | LAMA2 | SFRP1 | LAMC1 | APOA4* | GM20547* | MDH1 |
| COL12A1 | LAMB2 | SFRP2 | LUM | APOH* | GPD1 | MST1* |
| COL16A1 | LGALS3BP | SLIT2 | NID1 | AZGP1* | GPLD1* | MUG1* |
| COL1A1 | LGMN | SLIT3 | OGN | BTD* | GPX3 | PCCA |
| COL1A2 | LOX | SOD3 | PCOLCE | C1S2* | GSTM1 | PCX |
| COL4A1 | LOXL1 | SPARC | PEDF* | C4B* | H2AC20 | PGAM2 |
| COL4A2 | LOXL2 | SPARCL1 | POSTN | C4BPA* | H2BC3 | PGM1 |
| COL4A5 | LOXL3 | SPON2 | PXDN | C5* | H2-Q10* | PKM |
| COL5A2 | LPL | SRPX2 | QSOX1* | C6 | H4C1 | PLG* |
| COL5A3 | LTBP1 | SVEP1 | SERPINA3N* | C8A* | HBA-A1 | PPIB |
| COL6A1 | LTBP2 | TCN2 | | C8B* | HBB-B2 | PRDX1 |
| COL6A3 | LTBP3 | TGFBI | | C8G* | HBB-BS | PRDX2 |
| COL8A1 | LTBP4 | THBS1 | | C9* | HEP2* | PYGM |
| CPE | MATN2 | THBS2 | | CA2* | HGFAC* | PZP* |
| CRLF1 | METRNL | TIMP1 | | CES1B* | HP* | SERPINA3K* |
| CSF1 | MFAP5 | TIMP2 | | CES1C* | HPX* | SERPINA3M |
| CST3 | MFGE8 | TIMP3 | | CFHR1* | HRG* | TF* |
| CTGF | MGP | TNC | | CFHR4* | HSP90AA1 | TTR |
| CTSA | MMP2 | TSKU | | CFI | HSP90AB1 | TUBA1B |
| CTSB | MMP3 | WISP2 | | CKM | HSPA8 | TUBB4B |
| CTSL | NGF | ZFP462 | | CLU | ICA* | UBC |
| DAG1 | NID2 | | | COL15A1 | IGHG1 | VCP |
| DNAH7C | NTN4 | | | CPB2* | IGHG2C | VIM |
| DNAJB11 | NUCB1 | | | CPG* | IGHM | VTN |
| DPT | NUCB2 | | | CSRP3 | IGKC | 1700009N14RIK |
| ECM1 | OAF | | | EEF1A1 | IL1RAP* | |
| EFEMP1 | OLFML3 | | | EEF2 | ITIH1* | |

### Supplemental Data

**Supplemental Data 1: Secretome LC-MS/MS data.**

**Supplemental Data 2: Proteome LC-MS/MS data.**

**Supplemental Data 3: Transcriptome (SLAMseq) data.**

**Supplemental Data 4: In-vivo-secretome LC-MS/MS data.**
